# PET imaging of an antisense oligonucleotide in the living non-human primate brain using click chemistry

**DOI:** 10.1101/2025.02.06.636694

**Authors:** Brendon E. Cook, Thomas C. Pickel, Sangram Nag, Philippe N. Bolduc, Rouaa Beshr, Anton Forsberg Morén, Cathy Muste, Giulia Boscutti, Di Jiang, Long Yuan, Prodip Datta, Piotr Ochniewicz, Yasir Khani Meynaq, Sac-Pham Tang, Christophe Plisson, Mario Amatruda, Qize Zhang, Jonathan M. DuBois, Armin Delavari, Stephanie K. Klein, Ildiko Polyak, Adebowale Shoroye, Sara Girmay, Christer Halldin, Laurent Martarello, Emily A. Peterson, Maciej Kaliszczak

**Author notes:** These authors contributed equally to this work (Brendon E. Cook, Thomas C. Pickel).

## Abstract

Determination of a drug’s biodistribution is critical to ensure it reaches the target tissue of interest. This is particularly challenging in the brain where invasive sampling methods may not be possible. Here, a pretargeted imaging methodology is disclosed that utilizes bioorthogonal click chemistry to determine the distribution of an antisense oligonucleotide in the living brain following intrathecal dosing. A novel PET tracer, [^18^F]BIO-687, bearing a click-reactive *trans*-cyclooctene (TCO) was discovered and tested in conjunction with a Malat1 antisense oligonucleotide (ASO) conjugated with a methyltetrazine (MeTz). PET imaging in rats demonstrated that the tracer possesses good kinetic properties for CNS imaging and can react to form a covalent linkage with high specificity to the MeTz-conjugated-ASO *in vivo*. Further, the amount of tracer reacted by cycloaddition with the Tz was determined to be dependent on the concentration of ASO-MeTz in tissue, as determined through comparison of the imaging signal with the LC-MS of the tissue homogenate. The system was evaluated in cynomolgus monkeys, with PET imaging showing favorable tracer kinetics and specific binding to the ASO *in vivo*. These results demonstrate that the tracer [^18^F]BIO-687 can image intrathecally-delivered ASO distribution in the brain, and future studies should leverage this technology to evaluate ASO distribution in human subjects to study distribution.

**One Sentence Summary:** Distribution of an intrathecally administered antisense oligonucleotide can be imaged using a pretargeted approach in the living brains of non-human primates.

## Introduction

Drug delivery to the central nervous system is a challenging process, complicated by natural barriers and active elimination pathways designed to keep foreign infiltrates from damaging the most crucial organ system in the body. Due to these protective mechanisms, small molecules are most readily adapted for CNS-penetration, however, there have been recent efforts aimed at discovering biologics and antisense oligonucleotides (ASOs) to treat CNS diseases, which represents a more difficult endeavor. Due to the large size and physicochemical properties of biologics and ASOs, substantial effort is required to ensure that these potential drugs can be delivered safely to the CNS and reach the areas of disease, often requiring direct dosing into the cerebrospinal fluid (CSF) by intrathecal (IT) lumbar puncture. This effort is further complicated by the delicate nature of the CNS, as traditional sampling methods that are used to determine pharmacokinetics and biodistribution for peripherally active drugs may not be safe or practical for sampling CNS tissues.

Nuclear imaging is a key technology in drug development, enabling the characterization of clinical candidates non-invasively in human with numerous examples of applications for small molecules and biologics (*1*). As discussed above, there is limited information about ASO distribution in the human body, particularly for ASOs developed for CNS diseases. By tagging ASOs with a radionuclide, the distribution and clearance can be tracked noninvasively following dosing, and ideally can provide quantitative data on ASO concentration in various structures in the brain (*2*).

Typically, molecular imaging with molecules like ASOs, which display slow clearance and distribution kinetics, requires careful matching to a radionuclide with a similar physical decay half-life. This results in overall higher radiation dosimetry to subjects, as dosing of longer-lived isotopes and longer tissues residence times ultimately result in higher radiation exposure (*3*).

Our group recently demonstrated pretargeted imaging as a means to image ASO distribution in the CNS of rats (*4*). This approach utilized the extraordinarily fast, selective, and bioorthogonal inverse electron demand Diels-Alder (IEDDA) reaction between a 1,2,4,5-tetrazine (Tz) and a *trans-*cyclooctene (TCO) to introduce the radiolabel after administration of the drug, covalently binding the two *in vivo*.

IEDDA pretargeted imaging was initially proposed as a method for imaging and therapy of cancer, leveraging the specificity and selectivity of antibodies without the limitations to dosimetry imposed by traditional protein labeling strategies (*5*). Most strategies focused on labeling the antibody or vector with a TCO and several novel tetrazine-based radiotracers were developed, incorporating metal chelators to bind isotopes such as Cu-64, In-111, Tc-99m, Ga-68, and [^18^F]AlF (*6–11*). More recently, there have been efforts to develop tetrazine tracers incorporating F-18 without the need for chelators, better enabling optimization for CNS-penetrant tracers (*12, 13*). Indeed, several groups have published recent attempts at pretargeted imaging of antibodies in the brain using radiolabeled tetrazines (*14–16*).

Imaging of ASO CNS distribution proceeds as follows in Fig. 1: A) ASO is tagged with a non-radioactive click-reactive handle and dosed intrathecally; B) the ASO is allowed to distribute along the spine and into the brain; C) at timepoints of interest, a CNS-penetrant radiotracer with a complementary click-handle is administered intravenously. The radiotracer crosses the blood-brain-barrier and undergoes an irreversible click reaction to the ASO, while any unreacted tracer clears the body. Dynamic positron emission tomography (PET) imaging captures quantitative data on the distribution of ASO in the brain and spine. Lastly, because only a very small percentage of ASO reacts with the tracer, subsequent pretargeting scans are possible, targeting the remaining unreacted ASO.

**Fig. 1.**
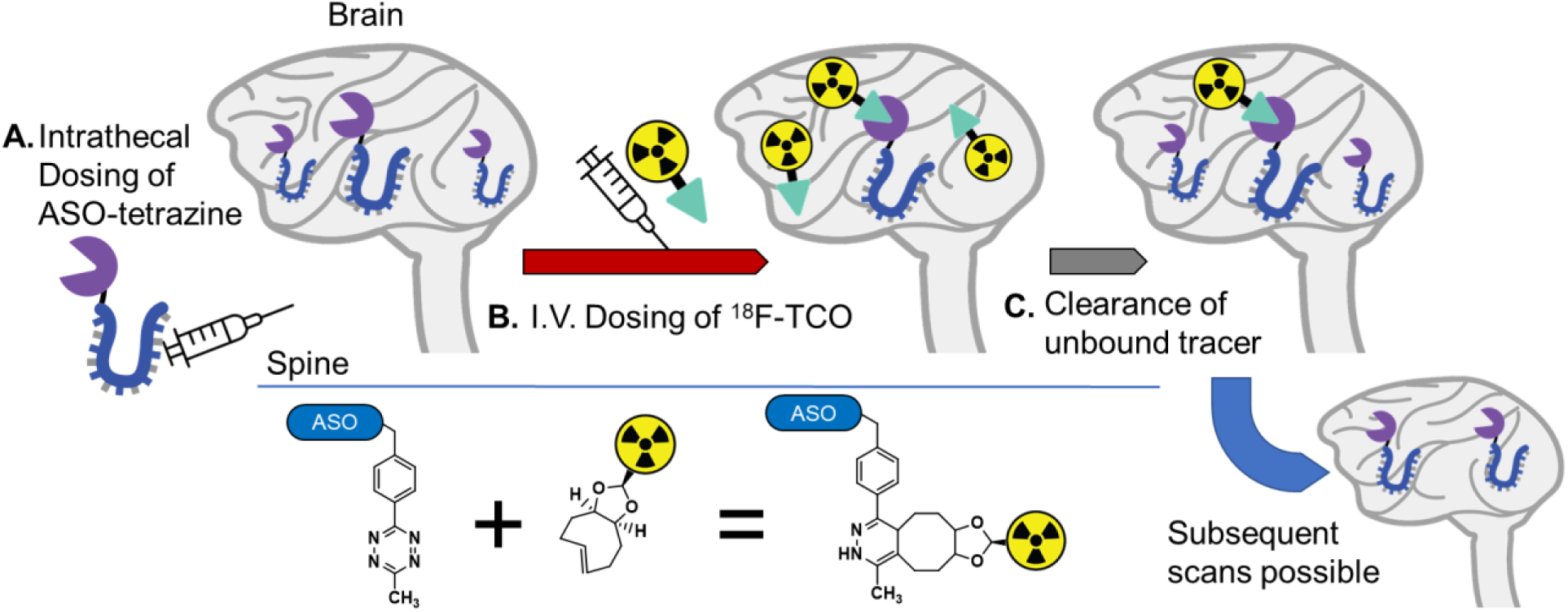
Schematic of pretargeted imaging. The IEDDA reaction between Tz and TCO is also shown below.

One of the key benefits of pretargeted imaging is the decoupling of the drug or vector of interest from the radioligand, enabling the use of shorter half-life radioisotopes, such as F-18 or C-11. This has been shown to drastically reduce the radiation dosimetry when compared to Zr-89 tagged molecules, potentially providing a safer, more translatable PET tracer (*6, 17*). This may also enable longer imaging timepoints that are not restricted by isotope decay. Longitudinal imaging to track the distribution of a single administration of ASO should be possible if the click reactive ASO is administered in molar excess relative to the PET tracer (*18*).

We previously demonstrated successful click reactions in the brain between a radiotracer [^18^F]F-537-Tz and Malat1 ASO-TCO, though the clearance of the tracer from the CNS was too slow to enable robust quantification and imaging in a non-human primate (NHP) (*4*). Presented here is a complete redesign of a pretargeted radiotracer capable of crossing the BBB and providing quantitative information on the distribution of antisense oligonucleotides in the CNS. Furthermore, evidence was generated in NHP demonstrating that this tracer will be suitable for human testing, potentially providing a first-in-class tool for the clinical imaging of ASOs by pretargeted imaging.

## Results

### Design of Click-Reactive ASO

Lessons learned from the previously disclosed approach led to a redesign of the ASO conjugate. It was disclosed previously that the TCO moiety had poor stability *in vivo*, with 50% isomerizing within 24 hours of dosing (*4*). Rather than continue to tag the ASO with a TCO, we instead tagged it with a methyltetrazine (MeTz) and designed the radiotracer to incorporate the TCO (*19, 20*).

In designing a tetrazine-bearing ASO, a critical consideration was to ensure stability of the click moiety. Three common tetrazines with different substitutions were selected, Me-Tz, H-Tz, and Pyr-Tz (Figure S1). Each of these was conjugated to the 5’ end of a Malat1 ASO via a hexylamino linker. These three constructs were then incubated in artificial cerebrospinal fluid (aCSF) for 72 hours at 37 °C and their stability checked by LC-MS (Figure S2). The Me-Tz ASO proved to be extremely stable with 97% parent remaining, while the H-Tz and Pyr-Tz samples showed lower stability with 74% and 20% parent remaining, respectively. These data are consistent with the known stability of the tetrazine moieties, themselves, with MeTz being highly stable, even in complex *in vitro* environments (*21*). These stability data were further supported *in vitro*, with ASO-MeTz showing the greatest stability after being incubated with HeLa cells in FBS-containing media for 24 hours, followed by staining with TCO-Cy5 dye (Figure S3).

The longitudinal *in vivo* stability of the Malat1 ASO-MeTz was tested next. Rats (n=5) were dosed intrathecally (IT) with 500 µg of Malat1 ASO-MeTz and then sacrificed at 1-, 2-, 5-, 7-, or 14-days post injection (n=1 per timepoint). Brains were removed, flash-frozen, sectioned, and stained for ASO and reactive ASO-MeTz (Fig. 2). In all animals, ASO and ASO-MeTz signal showed significant overlap, with the highest click-reactive signal at the 1-day timepoint. Immunofluorescence signal showed the highest uptake in what are likely perivascular cells, with more diffuse uptake seen in parenchymal cells.

**Fig. 2.**
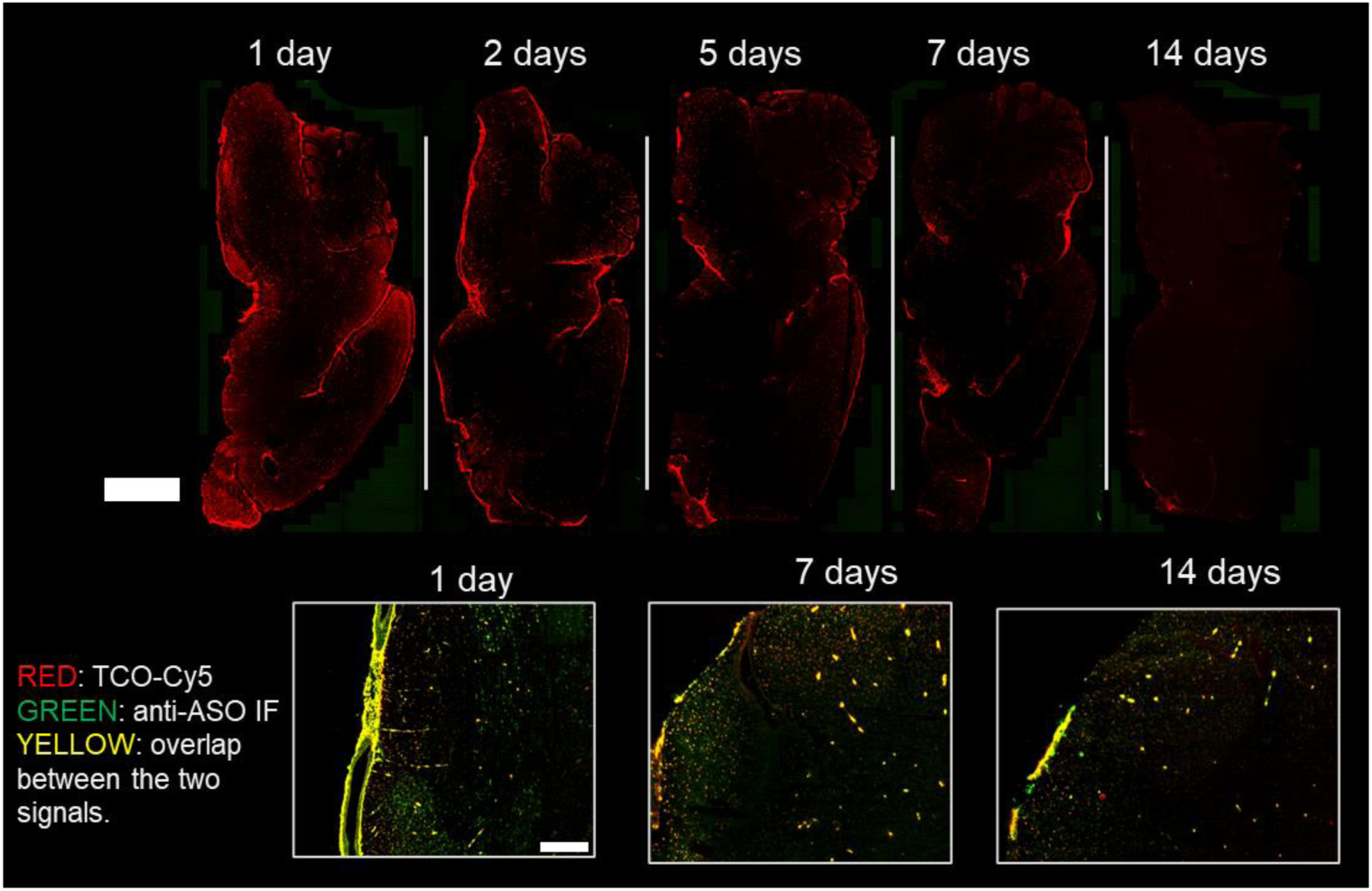
Immunofluorescence of ASO-MeTz in rats. IF images showing distribution of Malat1 ASO-MeTz in rat brains over 2-weeks after IT dosing. Top scale bar = 5 mm. Inserts show overlay of TCO-Cy5 (red), anti-ASO IF (green) and regions of overlap (yellow). Bottom scale bar = 200 µm.

Analysis of postmortem rat tissue by extraction and LC-MS was performed based on the novel protocol described in Materials and Methods. Using a TCO-PEG_4_-DBCO probe, the assay provides the concentration of both total Malat1 ASO-MeTz and Malat1 ASO-MeTz that remains reactive to TCO (Figure S4). This is a critical distinction, as tetrazine can undergo hydrolysis and reduction *in vivo*, resulting in very small changes to overall mass (<15 Da) that may not be detectable on traditional ASO mass spectrometry due to the complex tissue matrix present (*4*). This technique allows for the determination of the relative reactive-fraction of ASO-MeTz, or the percentage of ASO in tissue that remains reactive to TCO.

It was found that the ASO-MeTz signal decreased from 7.07 µg/mL at day 1 to 0.87 µg/mL at day 7, and 0.10 µg/mL at day 14 (Fig. 3). Notably, the concentration of ASO-MeTz (which includes non-reactive species) remained essentially identical to the concentration of ASO-MeTz-TCO-PEG_4_-DBCO, indicating that there was no measurable decomposition of the MeTz. Any instability in the compound is likely due to some loss of the MeTz or linker altogether rather than hydrolysis or reduction of the tetrazine.

**Fig. 3.**
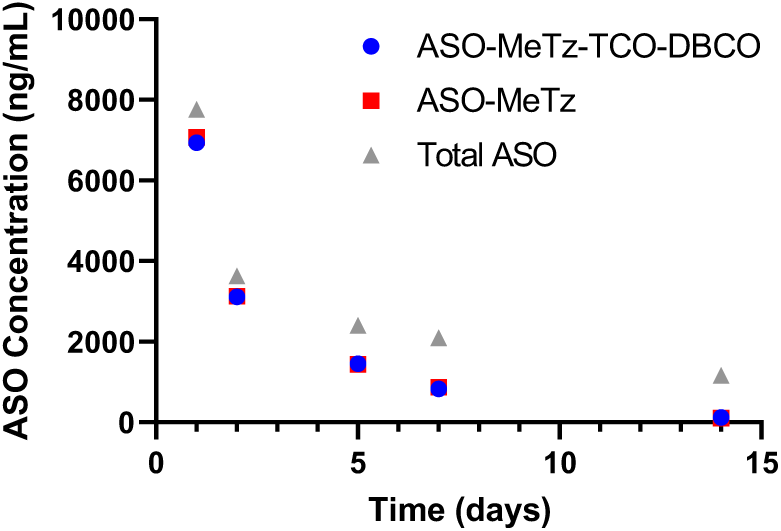
Stability of ASO-MeTz. Longitudinal stability and clearance of Malat1 ASO-MeTz in whole rat brain after IT dosing as measured by LC-MS. Graph shows total ASO concentration (grey triangle) which includes parent compound and any degradation products where linker or tetrazine was affected, ASO-MeTz (red square) where linker and tetrazine mass was unchanged, and ASO-MeTz-TCO-DBCO (blue circle) in which ASO-MeTz was reactive to a TCO-DBCO ligand.

### Design of a Trans-Cyclooctene (TCO) PET Tracer

With a suitable ASO-MeTz conjugate in hand, the identification of a complementary CNS-penetrant TCO radiotracer was investigated. PET ligand discovery, much like drug discovery, typically requires understanding the structure-activity relationship (SAR), minimizing off-target binding, and optimizing ADME properties. In the case of a TCO tracer, selectivity and click-reactivity with the Tz are almost exclusively a function of the *trans*-cyclooctene moiety and are not significantly impacted by other elements of the compound’s structure. Consequently, the focus of the optimization was the physicochemical properties on the portion of the molecule containing the radiolabel. The goal of this optimization was to enable a suitable biodistribution profile and facile radiolabel synthesis while also incorporating a UV handle for ease of detection during synthesis and analysis. Drawing on previously established guidelines for physicochemical property space of CNS penetrant drugs and combining that with ideal properties for PET tracers, key pharmacokinetic and physicochemical parameters were established for prospective TCO tracers (Table S1) (*22, 23*). Specifically, compounds were designed to have a CNS MPO score of >3, and properties that would result in low non-specific binding (>10% free in human and rat plasma and >5% in rat brain), a low MDR1 efflux ratio (< 3 in MDCK-BCRP and MDR1 assays), and rapid clearance (>50 mL/min/kg in human microsomes and >100 mL/min/kg in rat microsomes).

Prior to beginning discovery efforts, a survey of radiolabeled TCOs disclosed in the literature was conducted. To date, most of the reported labeled TCOs have been prepared by fluorination of a pendant ethoxy or PEG moiety(*24–26*). In the context of CNS PET ligand development, these functionalities present a liability owing to the potential formation of recirculating CNS penetrant radiolabeled metabolites (*27*). One reported tracer, ^18^F-fluoronicotinamide dioxolane TCO (dTCO) MICA-213 (**1**) (Fig. 4A), showed good uptake and clearance from the brain of rodents, and was used successfully in the pretargeted imaging of a peripheral tumor in conjunction with a tetrazine-antibody conjugate (*19*). Seeking to characterize this compound further, the non-radioactive MICA-213 was prepared and subjected to plasma protein binding (PPB) and microsomal clearance assays. MICA-213 was highly free in plasma (fraction unbound (Fu) = 28%, human; 52%, rodent) and brain tissue (Fu = 22%, rodent) and was rapidly metabolized by rat liver microsomes (264 mL/min/kg) but was significantly more long-lived in human liver microsomes (29 mL/min/kg). Despite these promising data, additional characterization of this compound was complicated by the propensity of the delicate TCO moiety to decompose quickly under ambient conditions.

**Fig. 4.**
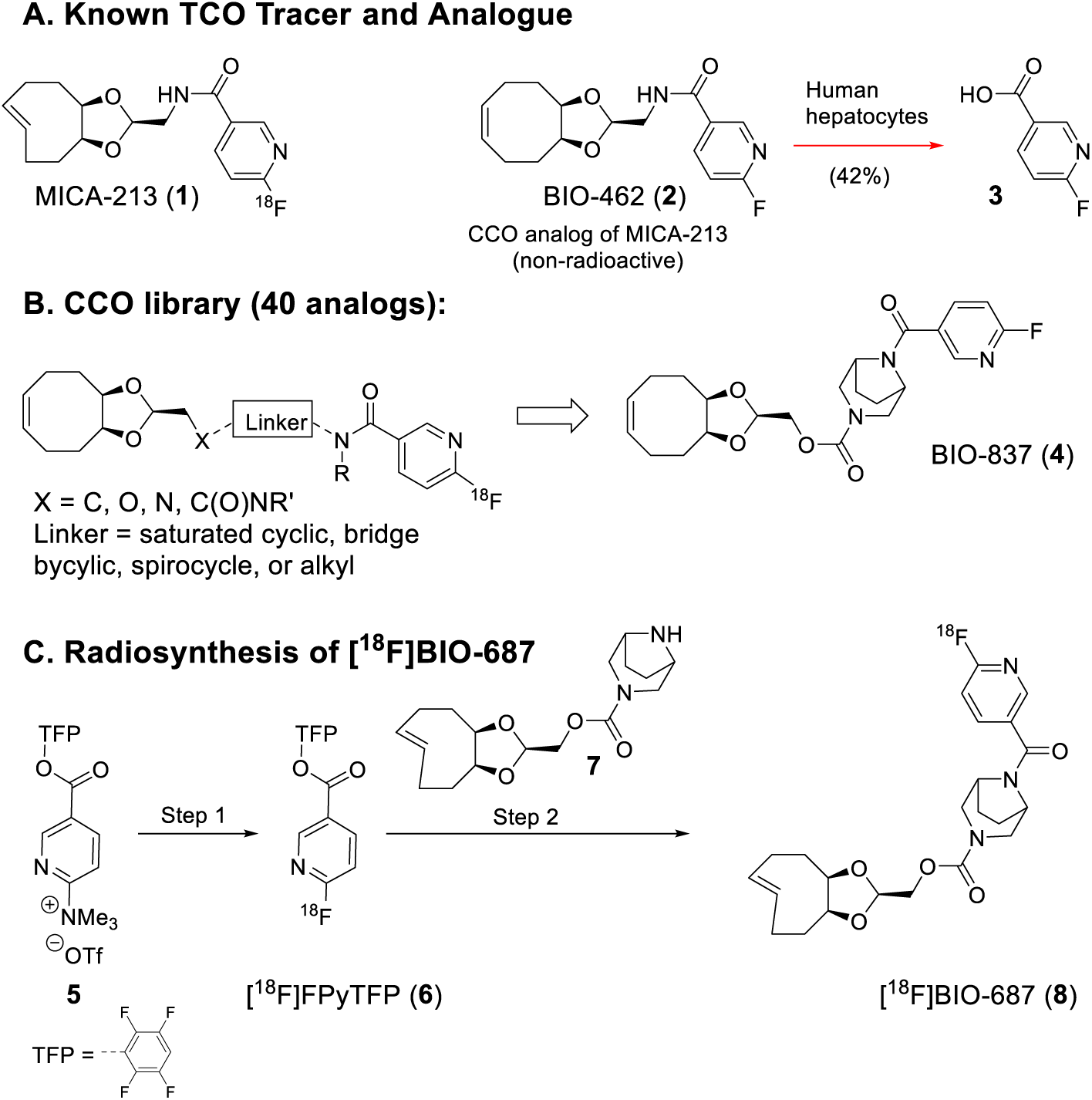
Screening, selection, and radiosynthesis of BIO-687. A) Reported TCO tracer MICA-213 and the CCO analog used to determine major human metabolites; B) Library of CCO analogs aimed to reduce amide hydrolysis risk but maintain CNS-penetration and optimal PET tracer properties; C) Radiosynthesis of [^18^F]BIO-687: Step 1: i. [^18^F]F^-^, K_2_CO_3_, K_2.2.2_ ii. Drying; iii. *t*-BuOH / MeCN (8:2), 80°C, 10 minutes; iv. SPE purification, elution in MeCN; Step 2: i. **7**, MeCN, 37 °C, 10 min; ii. HPLC purification, cartridge reformulation, and sterile filtration.

To circumvent TCO related stability challenges, the *cis*-cyclooctene (CCO) analogue of MICA-213 was prepared and characterized, with the hypothesis that the physicochemical and pharmacokinetic properties of the two compounds would be similar. Indeed, the fluoronicotinamide dCCO (BIO-462, **2**) gave comparable results in PPB (Fu = 23%, human; 44%, rodent) and brain tissue (Fu = 27%, rodent) assays as well as in *in vitro* microsomal clearance (334 mL/min/kg, rat; 31 mL/min/kg, human) assays (Table S1). Additionally, BIO-462 (**2**) afforded an efflux ratio of 0.71 in the MDCK-MDR1 permeability assay and a Kp_u,u_ of 0.49 (rat, 5 minutes post injection), indicating excellent brain uptake. Finally, the likelihood of CNS penetrant radiolabeled metabolite formation from BIO-462 (**2**) was evaluated by *in vitro* MET ID with rat, NHP, and human hepatocytes (Fig. S5). In lower species, all detectable metabolites were large and highly polar species produced by oxidation of the cyclooctene moiety and are unlikely to be highly CNS-penetrant. Conversely, in human hepatocytes, hydrolysis of the amide bond to form 6-fluoronicotinic acid (**3**) (a fluorinated analogue of niacin) was the major metabolite detected (42%) creating concerns that the radiolabeled portion of the tracer would be cleaved from the TCO click handle and form a brain penetrant radiolabeled metabolite (*28, 29*).

In view of these data, it was apparent that the structure of BIO-462 (**2**) required further optimization to improve the overall rate of metabolic clearance in human and prevent the observed amide hydrolysis while also minimizing formation of CNS permeable fluorinated metabolites. To enable the rapid identification of a tracer with the desired properties, the *cis*-cyclooctenes were utilized as surrogate TCOs. As such, a small library of CCO compounds with similar structural features to BIO-462 was prepared. The design of this library focused on diamine linkers that represented a range of stereoelectronic and steric features that could potentially reduce the amide cleavage metabolite and be paired with readily accessible derivatives of 5-OH and dTCO as well as electron deficient heteroarenes to promote relatively gentle ^18^F-fluoride labeling reactions (Fig. 4B) (*30, 31*). Each library compound was subjected to the previously described *in vitro* PPB, clearance, and efflux assays, culminating in the identification of *cis*-cyclooctene BIO-837 (**4**).

This compound was modestly effluxed by MDR1-MDCK cells (ER: 3.4) and displayed high free fraction in plasma (23%, human; 30%, rodent) and brain tissue (51%, rodent). Critically, BIO-837 (**4**) was rapidly metabolized in human and rat liver microsome assays (150 mL/min/kg and 826 mL/min/kg, respectively), and displayed excellent uptake and clearance from rat brain (Kp_u,u_ = 0.7, 5 min. post injection; not detectable in brain two hours post injection). In contrast to BIO-462 (**2**), metabolism of BIO-837 (**4**) occurred exclusively by oxidation of the cyclooctene ring in rodent, NHP, and human hepatocytes, and low molecular weight metabolites or those stemming from loss of fluoride were not detectable (Fig. S6).

Having identified a promising *cis*-cyclooctene analogue, a mild synthetic route to the *trans* analogue BIO-687 (**8**) was developed that would enable the incorporation of ^18^F to the delicate TCO moiety. Starting from TFP-protected ester pyridine (**5**), displacement of the quaternary amine with ^18^F was preferred over halogen precursors, which were harder to separate from the final product if unreacted (*32*). Displacement of the TFP-ester contained in radiolabeled ^18^FPyTFP (**6**) with dTCO amine (**7**) provided [^18^F]BIO-687 (**8**). With this synthetic approach, the dTCO moiety can be labeled quickly at room temperature in neutral media, reducing the likelihood of isomerization or oligomerization that is well established to occur at the elevated temperatures typically employed during the incorporation of ^18^F-fluoride.

### Radiochemistry

The radiolabeling of [^18^F]BIO-687 was achieved by production and purification of the amine-reactive prosthetic group [^18^F]FPyTFP, followed by coupling to the dTCO precursor in a second reaction vessel using a fully-automated procedure (Fig. 4C). After HPLC purification and SPE reformulation >1.0 GBq of [^18^F]BIO-687 was obtained from 15 min irradiation of the target at 35 µA. The average radiochemical yield (RCY) of the optimized radiosynthesis was 7.4 ± 4.3% (n=6, non-decay corrected), and the radiochemical purity (RCP) was >99%. The overall radiosynthesis including the fluorination reaction, HPLC purification, SPE isolation and radiotracer formulation was completed within 100-110 minutes. The identity of the labeled compound was confirmed by co-injection of [^18^F]BIO-687 and its corresponding fluorine-19 analogue using analytical HPLC (Fig. S7). The radioligand [^18^F]BIO-687 was found to be stable in formulation solution (pH=7.4) for at least 60 minutes.

Labeling was carried out in London for rodent studies and Stockholm for NHP studies. Rodent studies were completed first, when the radiosynthesis conditions were not yet fully optimized. Small changes in reaction time, temperature, and addition of a stabilizer (tocoferol) together with a new formulation resulted in higher RCY, RCP, and reproducibility of the radiotracer, which were then leveraged for studies in NHP. The fully optimized protocol is presented here, and results from all radiosyntheses used in rodent and NHP studies are summarized in Fig. S8.

### In Vitro Autoradiography

With both the radiotracer and ASO conjugate developed, an autoradiography study was run to ensure the two components could undergo a click reaction within a biological matrix. Rats were dosed i.t. with 500 µg of Malat1 ASO-MeTz and sacrificed 24 hours later. Brain sections were then incubated in a solution containing either [^18^F]BIO-687 tracer or tracer with 10 µM of the non-radioactive BIO-687. After washing and developing on a phosphor plate, the distribution of signal shows high uptake on the periphery and meninges of the brain, with a slow diffusion towards the midbrain (Fig. 5). This matches well with anti-ASO staining immunofluorescence on nearby sections. Furthermore, this binding signal is eliminated when tissue is incubated with a solution containing a homologous block, as well as when tracer is incubated with brain sections from a naïve rat. Overall, this demonstrates that binding of the tracer is highly specific to the presence of ASO-MeTz.

**Fig. 5.**
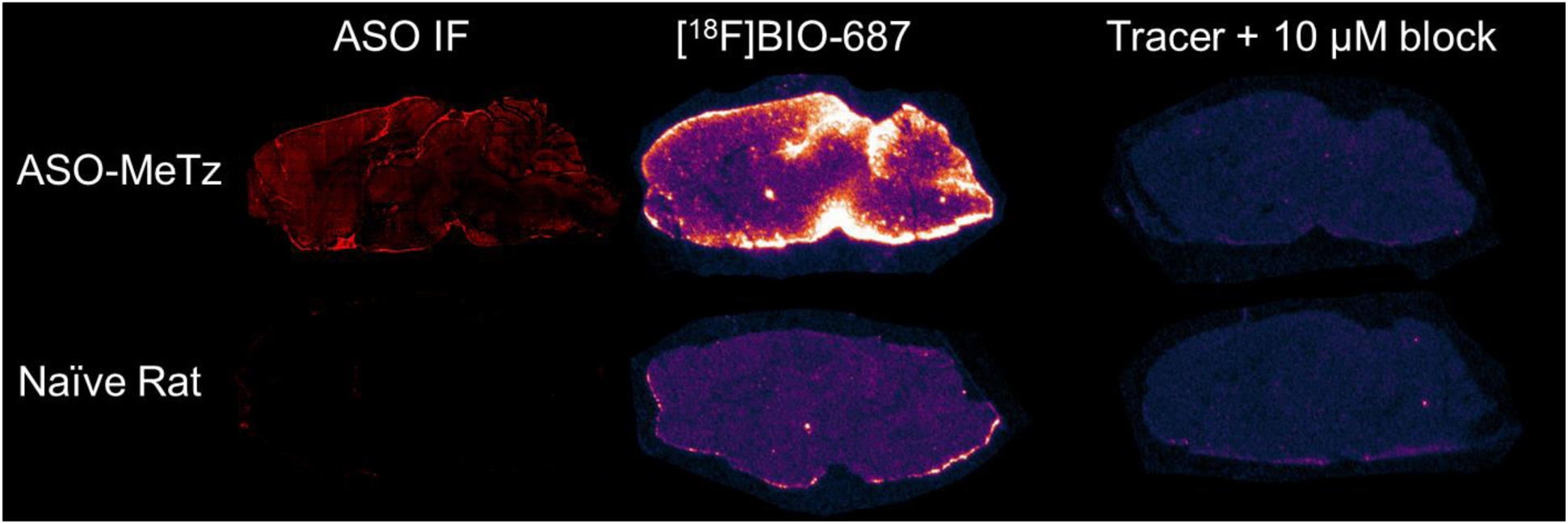
Autoradiography and IF. Rat dosed IT with 500 µg of Malat1 ASO-MeTz vs. naïve rat. Sacrificed 24 hours after dosing. Left to right, anti-ASO immunofluorescence, in vitro autoradiography with [^18^F]BIO-687, and block using 10 µM BIO-687.

### Pretargeted PET Imaging in Rat

The tracer [^18^F]BIO-687 was dosed in naïve rats (n=2) to characterize its baseline distribution and kinetics (Fig. S9). These rats received 60-minute dynamic PET scans following i.v. dosing of the tracer. Arterial blood sampling at prescribed timepoints was performed by in-dwelling catheter, allowing for determination of plasma parent fraction concentration. Brain uptake was observed with a C_max_ of ∼2.5 SUV, followed by rapid washout of >95% of signal by 20 minutes p.i. Parent fraction analysis showed intact tracer was present throughout the duration of the scan.

With *in vivo* behavior of the tracer showing favorable kinetics for PET imaging, it was next tested in rats following dosing of ASO. Three cohorts received Malat1 ASO-MeTz by i.t. injection: 750 µg, 500 µg, and 250 µg. A fourth cohort served as a “baseline” control, receiving 750 µg of authentic Malat1 ASO (not bioconjugated with tetrazine). After 24 h, each rat was scanned dynamically for 20 minutes upon the administration of a bolus i.v. dose of [^18^F]BIO-687. Immediately following the scan, blood and plasma samples were taken and counted by gamma counter, while brains, spine regions (lumbar, thoracic, cervical), and select peripheral organs (liver and kidneys) were removed for analysis by mass spectrometry to determine ASO concentration. Brains were dissected to isolate the frontal cortex, striatum, hippocampus, thalamus + hypothalamus, and cerebellum.

PET images show clear difference in tracer distribution in the CNS between pretargeted and baseline subjects (Fig. 6A). Uptake in the brain of animals receiving ASO-MeTz is characterized by higher uptake in the cortex with little uptake in the mid-brain, matching the pattern seen in the earlier autoradiography study (Fig. 5).

**Fig. 6.**
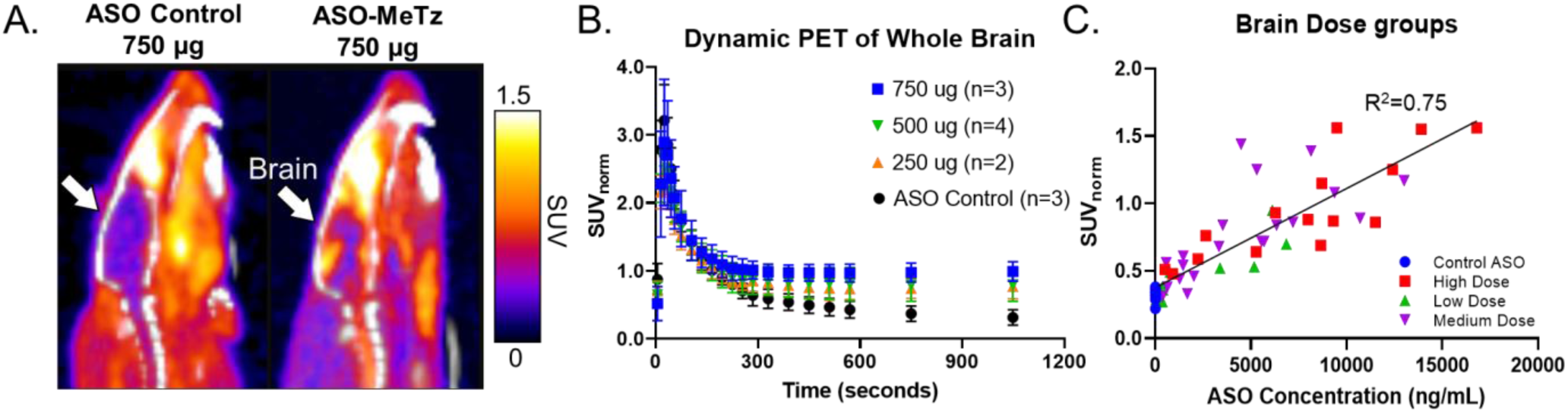
Pretargeted PET of ASO-MeTz in rat with [^18^F]BIO-687. Pretargeted PET imaging in rats dosed IT with 250-750 µg Malat1 ASO-MeTz; A) PET/CT images summed 0-20 min p.i. with [^18^F]BIO-687; B) time-activity curves; C) graph showing correlation between ASO concentration in brain subregions determined by LC-MS and PET SUV normalized to blood serum concentration of tracer; D) normalized SUV at the 20 minute timepoint.

Time-activity curves show the uptake of tracer in the brain throughout the duration of the scan (Fig. 6B). Tracer uptake determined by PET was normalized by dividing the SUV in brain tissue by SUV concentration of tracer in whole blood at 20 minutes p.i. (SUV_norm_). The control cohort shows the same uptake and washout seen in the baseline scans, with SUV_norm_ of ∼0.3 after 20 minutes, while the 750 µg pretargeted cohort stabilizes at ∼1.0 SUV_norm_. It is important to note that, since the ASO distribution and PET signal is highest around the edges of the brain and not specific to any subregions, the regions of interest (ROI) selected will dilute the overall signal. This will likely need to be addressed to properly interpret findings related to ASO distribution.

During post-mortem analysis, three animals (two in the 750 ug and one in the 250 ug cohort) were found to have brain ASO-MeTz concentrations below the limit of quantitation. These subjects were excluded from analysis, but their time-activity curves were mapped in Fig. S10. These curves were identical to the no-tetrazine ASO control cohort, providing further evidence that uptake of [^18^F]BIO-687 is dependent on the presence of ASO-MeTz in the CNS tissue.

Analysis of postmortem rat tissue by extraction and LC-MS was performed as described above to determine the concentration of ASO-MeTz, as well as the fraction still reactive to TCO (Fig. S11). Throughout the brain and spinal cord samples, the reactive fraction ranged from 77.4% in the striatum to 93.1% in the lumbar spinal cord (Fig. S12).

Bioanalysis of these samples also enabled a comparison between uptake of PET tracer and absolute quantified concentration of ASO in different brain subregions (Fig. 6C). Plotting the PET signal as SUV_norm_ against the concentration of ASO-MeTz in ng/mL shows a robust correlation (R^2^=0.75). It is important to note that the small size of the rat brain may lead to some discrepancy between digital ROIs and the dissected subregions, so it is likely that this correlation may be higher. Plotting values for the high, medium, and low doses of ASO demonstrates that, while there is variability in delivery within the different dose groups, overall trends show that a higher dose leads to higher tissue concentrations in the brain.

### Pretargeted PET Imaging in Non-Human Primates

With promising imaging results in rodents, the tracer [^18^F]BIO-687 was then tested in non-human primates (NHP). It was important to confirm that the tracer kinetics and brain uptake observed in rat would be similar in NHP. To this end, two NHPs (NHP1 and NHP2) were dosed [^18^F]BIO-687 by i.v. bolus and imaged by dynamic PET from 0-120 minutes p.i. (Fig. 7). Whole brain time-activity curves show rapid uptake in the brain followed by washout over the duration of the scan, with ∼90% clearance by 60-minutes p.i. (<0.5 SUV). Arterial blood samples were taken at discrete timepoints throughout the duration of the scan and analyzed for percent parent fraction by radioHPLC (Fig. S13). The parent tracer metabolizes quickly *in vivo*, with ∼30% remaining at 15 minutes p.i. Importantly, the two major radiometabolites observed elute before the parent compounds, indicating increased polarity and a low likelihood of entering the brain to confound PET signal.

**Fig. 7.**
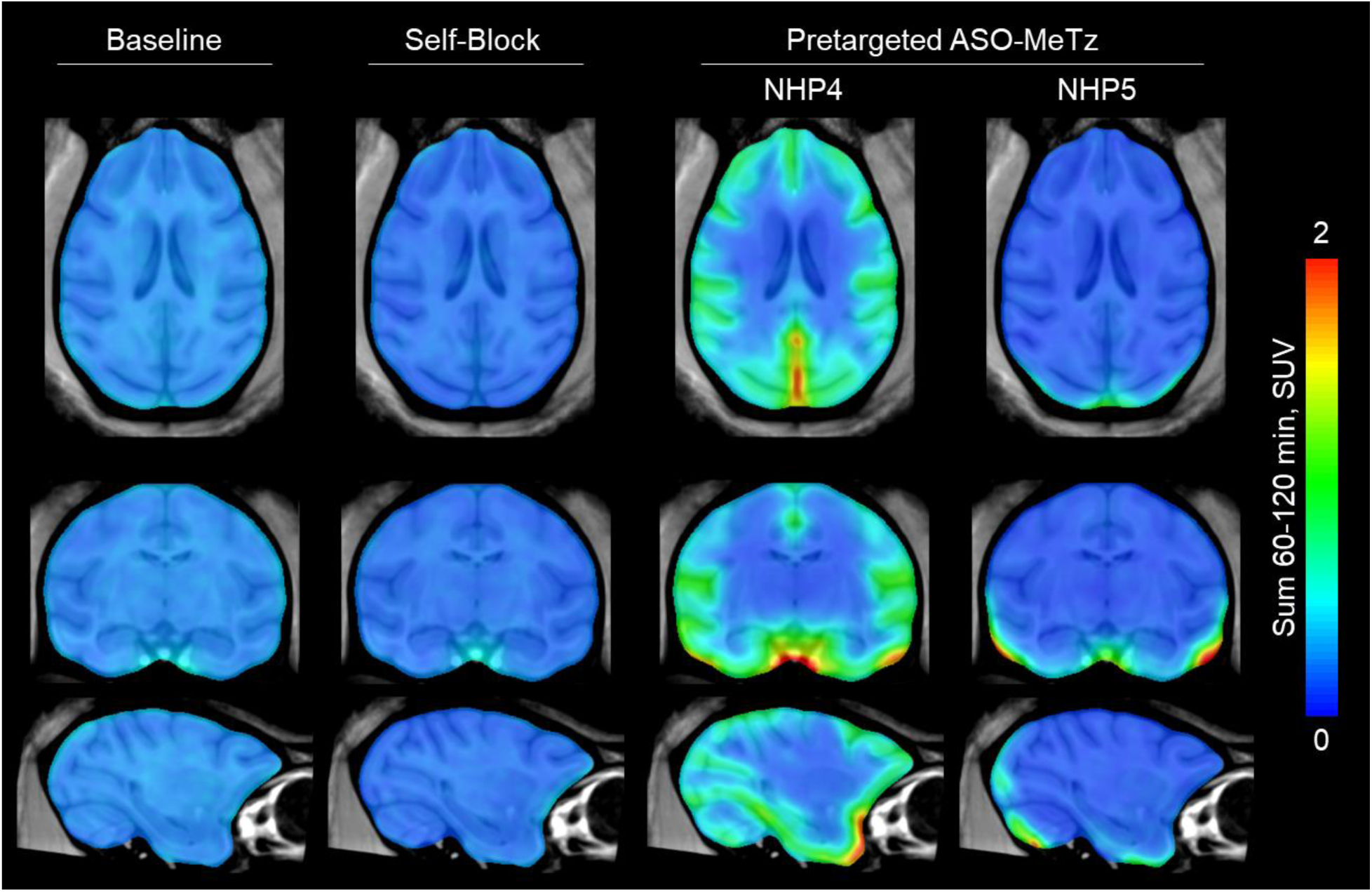
Pretargeted PET imaging in NHP. Left to right, representative baseline scan with [^18^F]BIO-687, representative non-radioactive self-block with 1.0 mg/kg BIO-687, pretargeted with 20 mg Malat1 ASO-MeTz dosed IT followed 24 hours later with tracer (NHP4 and NHP5). Images are summed from 60-120 minutes post-injection. The static SUV images, in a range of 0-2, are overlaid on a standard template T1-weighted MRI of the cynomolgus monkey, with a transparency of 0.7. The SUV images are smoothed at 2 mm FWHM and masked to show only the values within the brain. Unmasked images can be found in Fig. S16.

While there was very little signal retained in the brain, the lack of specific binding in the absence of ASO-MeTz was confirmed with a self-blocking experiment on two additional NHPs. NHPs were dosed with 1 mg/kg BIO-687 as a self-block before injection of the tracer [^18^F]BIO-687. There was no observed change in tracer kinetics following self-block, indicating that there is no detectable off-target binding in the brain (Fig. S14).

With excellent tracer kinetics and no observable saturable off-target binding in the brain in NHP, a pretargeted PET imaging study was performed following administration of Malat1 ASO-MeTz. The ASO-MeTz was dosed via intrathecal lumbar puncture (20 mg in 2.4 mL aCSF) into two NHPs. 24 hours after ASO dosing, [^18^F]BIO-687 was administered by i.v. bolus injection and the monkeys scanned by dynamic PET for 120 minutes. As before, arterial or venous blood was taken and analyzed for percent parent fraction of tracer and for the presence of radiometabolites (Fig. S14).

Higher uptake of PET tracer in the brain was observed in both animals that received doses of ASO-MeTz (Fig. 7). Critically, the distribution of PET signal is located primarily in cortical grey matter, which is expected given that the ASO was delivered intrathecally and is very similar to the observed distribution in rats. There were differences in brain signal between the two Malat1 ASO-MeTz dosed NHPs. NHP4 showed greater uptake throughout the cortex, with higher activity around the caudal area of the brain and near the spinal cord, consistent with ASO distribution following successful i.t. injection. NHP5, however, showed overall lower cortical distribution, though caudal accumulation of tracer was still present, as well as some activity in the hindbrain/cerebellum. Interestingly, it was noted during the IT dosing of ASO-MeTz that there was significant backflow of CSF following the lumbar puncture dosing, not seen during NHP4 dosing. It is quite possible that the backward pressure from the leaking CSF prevented full distribution of the ASO-MeTz to the brain of NHP5. It should also be noted that NHP5 weighed 7.0 kg, compared with NHP4 weighing 4.2 kg. This substantial difference in size could also be an important factor in the observed variability, potentially affecting spine length, diameter, and relative CSF flow.

Whole brain time-activity curves for the pretargeted subjects do not show significant separation from the baseline scan when the two subjects are averaged together. However, tracer clearance from NHP4 alone shows tracer retention, with 0.5 SUV at 120 minutes while baseline tracer is at 0.2 SUV (Fig.S14). This follows the expected pattern of an irreversible tracer binding its target and again mirrors what was seen in the rat imaging study. It should be noted that the use of standardized template regions does not adequately illustrate the true uptake of tracer, as ASO drug distribution is not homogeneous within each region. Particularly if pretargeted imaging will be used to estimate pharmacological dose delivered to the brain, it may be necessary to evaluate other non-traditional methods for analyzing the imaging data.

## Discussion

The ability to evaluate the tissue distribution and kinetics of ASOs, or any other therapeutic modality administered directly in the CNS, is critical to optimize dosing and to maximize the chance that drug exposure in the brain tissues most effected by the pathology reaches efficacious concentrations. Pre-targeted imaging as presented here could provide a means to accomplish this goal by using the novel TCO radiotracer, [^18^F]BIO-687, to measure the distribution of an ASO-tetrazine in the CNS.

While most groups developing pretargeting agents have selected tetrazines as the moieties for inclusion in the radiotracer, our previous experience demonstrated that this may not be the ideal choice for our applications. First, the TCO-conjugated ASO demonstrated instability by isomerization, precluding its use for imaging longer timepoints. Incorporating a MeTz into the ASO has the potential to greatly extend imaging once challenges around linker stability have been addressed. Second, it was found that TCO-based tracer candidates demonstrated more promising ADME properties and in vivo kinetics than Tz-based compounds. Furthermore, the reduced polar surface area of the TCO moiety when compared to Tz is more amenable to achieving CNS-penetration during SAR optimization.

Because the screened compounds exhibited robust membrane-crossing properties, the focus of structural optimization was on increasing rate of clearance while preventing BBB-penetrant radiometabolite formation. To this end, a bridged heterocycle linking the TCO to the radiolabeling moiety served well, producing rapid clearance from the plasma with only more-polar radiometabolites. It was demonstrated that a rapidly clearing tracer improves signal-to-noise by reducing non-specific background signal while still demonstrating suitable reactivity to achieve robust *in vivo* cycloaddition to the Tz-tagged target.

Initial imaging studies in rat provided key evidence supporting the functionality of the tracer [^18^F]BIO-687 for pretargeted imaging. PET images showed the distribution of the tracer was highest in CSF-proximal brain regions, agreeing well with ASO distribution that was seen in ex vivo immunofluorescence imaging and autoradiography. The specificity of the tracer for Malat1 ASO-MeTz was further supported by the LCMS enabled quantitation of ASO concentration in various subregions of the brain. The high degree of correlation between PET signal and absolute ASO concentration provides strong evidence that the tracer undergoes an irreversible click chemistry reaction in vivo, enabling non-invasive imaging and potential quantification of ASO distribution in living subjects.

Imaging studies in NHP established the translatability of this technique, demonstrating both the tracer’s superior kinetics in the CNS, as well as the ability for in vivo click chemistry to be performed in a larger primate species. Scans performed in NHP showed higher uptake in brain regions proximal to extra-cerebral CSF compartment after dosing with Malat1 ASO-MeTz when compared to baseline, again illustrating the ability of this tracer and methodology to evaluate the distribution of ASO *in vivo*.

One existing limitation of this system is the instability of the ASO-MeTz construct *in vivo*. Decomposition of the click-target over time adds an exceptional level of complexity to potential quantification efforts, as well as an inherent limitation to the timepoints that can be selected for imaging. While these data suggest that the tetrazine itself is not decomposing, further studies will be needed to assess the linkage between ASO and tetrazine to improve stability.

The development of drugs for applications in the CNS have encountered a growing number of roadblocks in recent years, often related to a lack of information on how well the drug can reach its intended target, or toxicity due to off-target distribution. With the evidence presented here, we believe pretargeted imaging will be an important tool to answer many of these questions, as it could be applied to imaging other therapeutic modalities in addition to ASOs. The modular nature of the TCO tracer and Tz handle allows for a low-profile, biocompatible modification to be applied to a wide range of macromolecules with minimal effect on biological activity (*4, 33, 34*).

We have demonstrated that pretargeting in the CNS can provide critical information on drug distribution following IT dosing and that PET signal is dependent on the concentration of tetrazine in tissue, suggesting that quantitative analysis may be possible. The PET tracer [^18^F]BIO-687 shows excellent uptake and clearance in the brain of cynomolgus monkeys, providing sufficient evidence to support clinical testing. If the observed brain kinetics are recapitulated in humans, this tracer could serve as a tool for testing clinical ASO distribution in the patients it is intended to treat.

## Materials and Methods

### Study Design

The goal of these studies was to design and test a novel PET tracer for pretargeted imaging that could image distribution of ASOs in non-human primates. Studies ranged from in vivo cell culture in HeLa cells, through initial proof-of-concept in rats, and finally imaging in non-human primates. Each individual study is described in more detail below.

Sample size for *in vivo* studies was determined with the goal of using the smallest number of animals that would still allow for robust interpretation of the data. The primary source of variability in these studies is the intrathecal dosing, and the number of animals dosed (n=3-4) was selected based on our previous experience with this method. Non-human primate studies were powered with n=2 per cohort to provide evidence of reproducibility, while also acknowledging the current scarcity of available NHPs for research.

### Synthesis of Malat1 ASO-Methyltetrazine

Synthesis procedure for generating Malat1 ASO tetrazine analogues is provided in the Supplemental Information.

### Synthesis and In Vitro Testing of Chemical Compounds

Detailed synthesis of BIO-462, BIO-687, BIO-837, and Compound 1 are provided in the Supplemental Information. (*32, 35*)

### Development of an LC-MS assay for detecting reactive Malat1 ASO-MeTz

Rat brains were weighed and homogenized in 9-fold (v/w) of lysis-loading buffer using a MP Biomedicals FastPrep-24 homogenizer, yielding 10-fold diluted tissue homogenate. A calibration curve and quality control (QC) samples of known Malat1 ASO and Malat1 ASO-MeTz concentrations were prepared in the tissue homogenate. The ASOs were extracted from the tissue, calibration curve, and QC samples via hybridization extraction using magnetic beads. After extraction, TCO-PEG4-DBCO was added to conjugate with Malat1 ASO-MeTz for the analysis of reactive Malat1 ASO-MeTz.

The concentrations of Malat1 ASO and reactive Malat1 ASO-MeTz were analyzed by LC-MS which included a Clarity Oligo-XT column (2.1 x 50 mm, 1.7 µm particle size) using mobile phases HFIP and DMCHA in water and acetonitrile with a gradient HPLC method (2–20 % in 8 min) and flow rate of 0.5 mL/min. MS detection was conducted using a Sciex QTRAP 6500+ mass spectrometer in negative ion mode using multiple reaction monitoring (MRM).

### Pretargeted PET Imaging in Rat

Naïve male Sprague Dawley rats (N = 15, 311 ± 41 g) were anesthetized under recovery anesthesia (isoflurane ca. 1.5 – 3%, 1 L oxygen·min-1) and a catheter inserted into the spinal canal for IT administration (30 μL dose followed by 40 μL saline flush) of ASO control or Malat1 ASO-MeTz as follows: high dose (750 μg) of Malat1 ASO control (N = 3), high dose (750 μg) of Malat1 ASO-MeTz (N = 5), low dose (250 μg) of Malat1 ASO-MeTz (N = 3), medium dose (500 μg) of Malat1 ASO-MeTz (N = 4).

24 h after IT administration, rats were placed under terminal anesthesia (isoflurane ca. 1.5-3%, 1 L oxygen.min-1) and a catheter was placed in a vein. Body temperature was monitored and maintained, and the respiration rate was monitored throughout the experiment. A 10-minute CT scan was collected for attenuation/scatter correction and structural information. This was followed immediately by a dynamic PET scan of 20 minutes, which was acquired on each of the rats after IV injection of [^18^F]BIO-687 (6.42 ± 1.10 MBq, 0.55 ± 0.36 µg), with a field of view focused on the brain and upper body.

Following the scans, animals were euthanized by exsanguination followed by cervical dislocation. Exsanguinated arterial blood sample was collected for radioactivity measurement at the end of the scan (in blood and plasma). Ten tissue samples were collected: brain regions (frontal cortex, striatum, hippocampus, thalamus+ hypothalamus, cerebellum), spinal cord (cervical, thoracic, lumbar), liver and kidneys. Tissue samples were placed into Lysing matrix D tubes, weighed, flash frozen in liquid nitrogen, stored at -80 °C until analysis by LC-MS.

Regions of interest in the PET scans were defined over 8 brain regions (frontal cortex, striatum, cortex, hypothalamus, thalamus, hippocampus, cerebellum, whole brain) as well as heart, liver, muscle, cervical and thoracic spine, to generate time-activity curves.

### Pretargeted PET Imaging in NHP

The study was approved by the Stockholm ethical committee (Stockholms djurförsöksetiska nämnd) of the Swedish Animal Welfare Agency (Dnr 10367-2019) and was performed according to “Guidelines for planning, conducting and documenting experimental research” of Karolinska Institutet.

For baseline experiments, two female cynomolgus non-human primates (NHPs) were used (NHP1, 5.2 kg; NHP2, 6.0 kg). The NHPs were housed in the Astrid Fagraeus laboratory (KM-F), Comparative Medicine Department at Karolinska Instituitet (Solna, Sweden). PET experiments were conducted using the MultiScan LFER 150 PET/CT system (Mediso Ltd.). Anesthesia was initiated by an intramuscular ketamine injection (10 mg/kg) and maintained by the administration of a mixture of sevoflurane, oxygen, and medical air after endotracheal intubation. Oxygen saturation, heart and respiratory rates, and blood pressure were continuously monitored all through the scan. Body temperature was maintained by a Bair Hugger-Model 505 (Arizant Healthcare Inc., MN) and monitored with an esophageal thermometer. The head was immobilized throughout scanning with a fixation device and fluid balance was maintained by a continuous infusion of Ringer Acetate. [^18^F]BIO-687 was injected as an intravenous bolus (158 MBq in NHP1, 152 MBq in NHP2) simultaneously with the start of PET data acquisition. Brain radioactivity was measured continuously for 125 minutes according to a preprogrammed series of 35 frames. For both NHPs, arterial blood sampling using a permanent catheter placed into the femoral artery, were performed at different time points for the measurement of blood and plasma radioactivity and radiometabolite HPLC analysis.

For experiments including IT administration, one female (NHP4, 4.2 kg) and one male (NHP5, 7.0 kg) NHP were used. The intrathecal administration of the Malat1 ASO-MeTz dosing was performed 24 h prior to [^18^F]BIO-687 administration. Following sedation with an intramuscular injection of ketamine (10 mg/kg), animals were positioned in abdominal recumbency, and a lumbar puncture was performed in the L4/L5 or L5/L6 intervertebral space using a 25G spinal needle with an introducer. The ASO-MeTz (20 mg per animal) was dissolved in the aCSF (Bio-Techne) and injected over 2-6 minutes in volume of 2.4 mL. The PET experiment was carried out as described above. [^18^F]BIO-687 was injected as an intravenous bolus (145 MBq in NHP4, 142 MBq in NHP5) simultaneously with the start of PET data acquisition. Venous blood sampling in NHP4 and arterial blood sampling in NHP5, using a permanent catheter placed into the femoral artery, were performed at different time points for the measurement of blood and plasma radioactivity and radiometabolite analysis.

Summed PET images of the whole PET acquisition were coregistered manually to the T1-weighted brain magnetic resonance (MR) image and regions of interest (ROI) were delineated manually for the whole brain, occipital cortex, caudate, putamen, ventral striatum, frontal cortex, white matter, thalamus, cerebellum, and hippocampus. The time-activity curves of brain regions were generated from dynamic PET data after the application of coregistration parameters to ROIs. Regional uptake was calculated as standardized uptake value (SUV), which is defined as uptake (Bq/mL)/injected radioactivity (Bq) × body weight (g).

#Fig. 7 underwent the following processing to best highlight tracer uptake and distribution within the brain. Dynamic frames were averaged from 60-120 minutes post-injection to create static volumes, and SUVs were calculated as described above. For each NHP, registration was then performed between the static SUV volume and the T1-weighted MRI using Advanced Neuroimaging Tools (*36*). In addition, each NHP MRI was registered to a template T1-weighted MRI generated from 18 cynomolgus monkeys (*37*). Each registration was visually inspected to confirm accurate alignment. The SUV data was then linearly resampled from native space to the template space and a Gaussian blurring kernel of 2 mm full width at half maximum (FWHM) was applied. Lastly, the template brain mask was used to mask the SUV images to show only the values within the brain. Fig. S16 displays brain PET as processed above but without application of the brain mask.

## List of Supplementary Materials

Supplementary Table S1

Supplementary Fig. S1 to S16

Materials and Methods

## Acknowledgements

The authors would like to thank the Biogen Cellular Imaging Unit (CIU) and the Biogen In Vivo Resource Center (IRC).

## Funding

All work in this paper was funded in full by Biogen.

## Competing Interests

Authors are/were employed or contracted by Biogen or Ionis and may hold stock in one or both companies. No other conflicts of interest exist.

## Data and materials availability

All data associated with this study are present in the paper or the Supplementary Materials. Sufficient methodology is provided to synthesize ASO conjugate and PET tracer.

## References

1. M. Honer, L. Gobbi, L. Martarello, R. A. Comley, Radioligand development for molecular imaging of the central nervous system with positron emission tomography. Drug Discovery Today 19, 1936–1944 (2014).

2. C. Mazur, B. Powers, K. Zasadny, J. M. Sullivan, H. Dimant, F. Kamme, J. Hesterman, J. Matson, M. Oestergaard, M. Seaman, R. W. Holt, M. Qutaish, I. Polyak, R. Coelho, V. Gottumukkala, C. M. Gaut, M. Berridge, N. J. Albargothy, L. Kelly, R. O. Carare, J. Hoppin, H. Kordasiewicz, E. E. Swayze, A. Verma, Brain pharmacology of intrathecal antisense oligonucleotides revealed through multimodal imaging. JCI Insight 4, (2019).

3. M. Monine, D. Norris, Y. Wang, I. Nestorov, A physiologically-based pharmacokinetic model to describe antisense oligonucleotide distribution after intrathecal administration. Journal of Pharmacokinetics and Pharmacodynamics 48, 639–654 (2021).

4. B. E. Cook, J. Archbold, K. Nasr, S. Girmay, S. I. Goldstein, P. Li, S. Dandapani, N. E. Genung, S.-P. Tang, S. McClusky, C. Plisson, M. E. Afetian, C. A. Dwyer, M. Fazio, W. J. Drury, F. Rigo, L. Martarello, M. Kaliszczak, Non-invasive Imaging of Antisense Oligonucleotides in the Brain via In Vivo Click Chemistry. Molecular Imaging and Biology 24, 940–949 (2022).

5. R. Rossin, P. Renart Verkerk, S. M. van den Bosch, R. C. M. Vulders, I. Verel, J. Lub, M. S. Robillard, In Vivo Chemistry for Pretargeted Tumor Imaging in Live Mice. Angewandte Chemie International Edition 49, 3375–3378 (2010).

6. B. M. Zeglis, C. Brand, D. Abdel-Atti, K. E. Carnazza, B. E. Cook, S. Carlin, T. Reiner, J. S. Lewis, Optimization of a Pretargeted Strategy for the PET Imaging of Colorectal Carcinoma via the Modulation of Radioligand Pharmacokinetics. Molecular Pharmaceutics 12, 3575–3587 (2015).

7. R. Rossin, S. M. van Duijnhoven, T. Läppchen, S. M. van den Bosch, M. S. Robillard, Trans-cyclooctene tag with improved properties for tumor pretargeting with the diels-alder reaction. Molecular Pharmaceutics 11, 3090–3096 (2014).

8. M. F. García, X. Zhang, M. Shah, J. Newton-Northup, P. Cabral, H. Cerecetto, T. Quinn, (99m)Tc-bioorthogonal click chemistry reagent for in vivo pretargeted imaging. Bioorganic & Medicinal Chemistry 24, 1209–1215 (2016).

9. J. P. Meyer, P. Kozlowski, J. Jackson, K. M. Cunanan, P. Adumeau, T. R. Dilling, B. M. Zeglis, J. S. Lewis, Exploring Structural Parameters for Pretargeting Radioligand Optimization. Journal of Medicinal Chemistry 60, 8201–8217 (2017).

10. C. A. Maitz, S. Delaney, B. E. Cook, A. R. Genady, R. Hoerres, M. Kuchuk, G. Makris, J. F. Valliant, S. Sadeghi, J. S. Lewis, H. M. Hennkens, J. N. Bryan, B. M. Zeglis, Pretargeted PET of Osteodestructive Lesions in Dogs. Molecular Pharmaceutics, (2022).

11. H. A. Bilton, Z. Ahmad, N. Janzen, S. Czorny, J. F. Valliant, Preparation and Evaluation of 99mTc-labeled Tridentate Chelates for Pre-targeting Using Bioorthogonal Chemistry. Journal of Visual Experiments, (2017).

12. E. J. L. Stéen, J. T. Jørgensen, C. Denk, U. M. Battisti, K. Nørregaard, P. E. Edem, K. Bratteby, V. Shalgunov, M. Wilkovitsch, D. Svatunek, C. B. M. Poulie, L. Hvass, M. Simón, T. Wanek, R. Rossin, M. Robillard, J. L. Kristensen, H. Mikula, A. Kjaer, M. M. Herth, Lipophilicity and Click Reactivity Determine the Performance of Bioorthogonal Tetrazine Tools in Pretargeted In Vivo Chemistry. ACS Pharmacology & Translational Science 4, 824–833 (2021).

13. U. M. Battisti, K. Bratteby, J. T. Jørgensen, L. Hvass, V. Shalgunov, H. Mikula, A. Kjær, M. M. Herth, Development of the First Aliphatic 18F-Labeled Tetrazine Suitable for Pretargeted PET Imaging—Expanding the Bioorthogonal Tool Box. Journal of Medicinal Chemistry 64, 15297–15312 (2021).

14. C. Bredack, M. R. Edelmann, E. Borroni, L. C. Gobbi, M. Honer, Antibody-Based In Vivo Imaging of Central Nervous System Targets-Evaluation of a Pretargeting Approach Utilizing a TCO-Conjugated Brain Shuttle Antibody and Radiolabeled Tetrazines. Pharmaceuticals (Basel) 15, (2022).

15. K. A. Morgan, M. de Veer, L. A. Miles, C. Kelderman, C. McLean, C. L. Masters, K. Barnham, J. White, B. M. Paterson, P. S. Donnelly, Pre-Targeting Amyloid-β with Antibodies for Potential Molecular Imaging of Alzheimer’s Disease. Chemical Communications, (2023).

16. S. Lopes van den Broek, V. Shalgunov, R. García Vázquez, N. Beschorner, N. S. R. Bidesi, M. Nedergaard, G. M. Knudsen, D. Sehlin, S. Syvänen, M. M. Herth, Pretargeted Imaging beyond the Blood-Brain Barrier: Utopia or Feasible? Pharmaceuticals 15, 1191 (2022).

17. R. Rossin, T. Läppchen, S. M. van den Bosch, R. Laforest, M. S. Robillard, Diels–Alder Reaction for Tumor Pretargeting: In Vivo Chemistry Can Boost Tumor Radiation Dose Compared with Directly Labeled Antibody. Journal of Nuclear Medicine 54, 1989 (2013).

18. O. Keinänen, J. M. Brennan, R. Membreno, K. Fung, K. Gangangari, E. J. Dayts, C. J. Williams, B. M. Zeglis, Dual Radionuclide Theranostic Pretargeting. Molecular Pharmaceutics 16, 4416–4421 (2019).

19. E. Ruivo, F. Elvas, K. Adhikari, C. Vangestel, G. Van Haesendonck, F. Lemière, S. Staelens, S. Stroobants, P. Van der Veken, L. wyffels, K. Augustyns, Preclinical Evaluation of a Novel 18F-Labeled dTCO-Amide Derivative for Bioorthogonal Pretargeted Positron Emission Tomography Imaging. ACS Omega 5, 4449–4456 (2020).

20. E. M. F. Billaud, S. Belderbos, F. Cleeren, W. Maes, M. Van de Wouwer, M. Koole, A. Verbruggen, U. Himmelreich, N. Geukens, G. Bormans, Pretargeted PET Imaging Using a Bioorthogonal 18F-Labeled trans-Cyclooctene in an Ovarian Carcinoma Model. Bioconjugate Chemistry 28, 2915–2920 (2017).

21. J. M. Mejia Oneto, I. Khan, L. Seebald, M. Royzen, In Vivo Bioorthogonal Chemistry Enables Local Hydrogel and Systemic Pro-Drug To Treat Soft Tissue Sarcoma. ACS Central Science 2, 476–482 (2016).

22. T. T. Wager, X. Hou, P. R. Verhoest, A. Villalobos, Central Nervous System Multiparameter Optimization Desirability: Application in Drug Discovery. ACS Chemical Neuroscience 7, 767–775 (2016).

23. L. Zhang, A. Villalobos, Strategies to facilitate the discovery of novel CNS PET ligands. EJNMMI Radiopharmaceutical Chemistry 1, 13 (2017).

24. M. Wang, R. Vannam, W. D. Lambert, Y. Xie, H. Wang, B. Giglio, X. Ma, Z. Wu, J. Fox, Z. Li, Hydrophilic 18F-labeled trans-5-oxocene (oxoTCO) for efficient construction of PET agents with improved tumor-to-background ratios in neurotensin receptor (NTR) imaging. Chemical Communications 55, 2485–2488 (2019).

25. L. wyffels, D. Thomae, A.-M. Waldron, J. Fissers, S. Dedeurwaerdere, P. Van der Veken, J. Joossens, S. Stroobants, K. Augustyns, S. Staelens, In vivo evaluation of 18F-labeled TCO for pre-targeted PET imaging in the brain. Nuclear Medicine and Biology 41, 513–523 (2014).

26. M. Wang, D. Svatunek, K. Rohlfing, Y. Liu, H. Wang, B. Giglio, H. Yuan, Z. Wu, Z. Li, J. Fox, Conformationally Strained trans-Cyclooctene (sTCO) Enables the Rapid Construction of (18)F-PET Probes via Tetrazine Ligation. Theranostics 6, 887–895 (2016).

27. V. W. Pike, PET radiotracers: crossing the blood-brain barrier and surviving metabolism. Trends in Pharmacological Sciences 30, 431–440 (2009).

28. L. V. Hankes, H. H. Coenen, E. Rota, K. J. Langen, H. Herzog, W. Wutz, G. Stoecklin, L. E. Feinendegen, Effect of Huntington’s and Alzheimer’s diseases on the transport of nicotinic acid or nicotinamide across the human blood-brain barrier. Advances in Experimental Medicine and Biology 294, 675–678 (1991).

29. J. Chen, M. Chopp, Niacin, an Old Drug, has New Effects on Central Nervous System Disease. Open Drug Discovery Journal 2, (2010).

30. M. Royzen, G. P. A. Yap, J. M. Fox, A Photochemical Synthesis of Functionalized trans-Cyclooctenes Driven by Metal Complexation. Journal of the American Chemical Society 130, 3760–3761 (2008).

31. A. Darko, S. Wallace, O. Dmitrenko, M. M. Machovina, R. A. Mehl, J. W. Chin, J. M. Fox, Conformationally strained trans-cyclooctene with improved stability and excellent reactivity in tetrazine ligation. Chemical Science 5, 3770–3776 (2014).

32. D. E. Olberg, J. M. Arukwe, D. Grace, O. K. Hjelstuen, M. Solbakken, G. M. Kindberg, A. Cuthbertson, One Step Radiosynthesis of 6-[18F]Fluoronicotinic Acid 2,3,5,6-Tetrafluorophenyl Ester ([18F]F-Py-TFP): A New Prosthetic Group for Efficient Labeling of Biomolecules with Fluorine-18. Journal of Medicinal Chemistry 53, 1732–1740 (2010).

33. J. W. Seo, E. S. Ingham, L. Mahakian, S. Tumbale, B. Wu, S. Aghevlian, S. Shams, M. Baikoghli, P. Jain, X. Ding, N. Goeden, T. Dobreva, N. C. Flytzanis, M. Chavez, K. Singhal, R. Leib, M. L. James, D. J. Segal, R. H. Cheng, E. A. Silva, V. Gradinaru, K. W. Ferrara, Positron emission tomography imaging of novel AAV capsids maps rapid brain accumulation. Nature Communications 11, 2102 (2020).

34. B. E. Cook, R. Membreno, B. M. Zeglis, Dendrimer Scaffold for the Amplification of In Vivo Pretargeting Ligations. Bioconjugate Chemistry 29, 2734–2740 (2018).

35. J. M. Fairhall, J. C. Camilli, B. H. Gibson, S. Hook, A. B. Gamble, EGFR-targeted prodrug activation using bioorthogonal alkene-azide click-and-release chemistry. Bioorganic & Medicinal Chemistry 46, 116361 (2021).

36. B. B. Avants, N. J. Tustison, G. Song, P. A. Cook, A. Klein, J. C. Gee, A reproducible evaluation of ANTs similarity metric performance in brain image registration. Neuroimage 54, 2033–2044 (2011).

37. S. Frey, D. N. Pandya, M. M. Chakravarty, L. Bailey, M. Petrides, D. L. Collins, An MRI based average macaque monkey stereotaxic atlas and space (MNI monkey space). NeuroImage 55, 1435–1442 (2011).

