## Supplementary Materials for "PET imaging of an antisense oligonucleotide in the living non-human primate brain using click chemistry"

Figure S15....... **Error! Bookmark not defined.**

### Supplemental Figures

#### Table S1

| **Specific Measure** | **PET Tracer Preferred Criteria*** | **BIO-462** | **BIO-837** (CCO) | **BIO-687** (TCO) |
|---|---|---|---|---|
| Brain Binding (R) % unbound | >5% | 27 | 51 | 16 |
| Plasma Protein Binding (H / R) % unbound | >10% | 23 / 44 | 23 / 30 | 32 / 45 |
| MDR1-MDCK ER [Papp AB (nm/s)] | < 3, [> 50 nm/s] | 0.7 [404] | 3.4 [116] | - |
| MDCK-BCRP ER | < 2 | ND | 0.56 | - |
| ELogD | 1.0 < Log D < 3.0 | 2.0 | 1.8 | - |
| HLM / RLM / CyLM Clint (mL/min/kg) | >50 / >100 | 31 / 334 / 936 | 150 / 826 / >1300 | - |
| Predicted T ½ (min) H/ R / Cy | < 2 h | 40 / 8 / 2 | 8.3 / 3.4 / 1.4 | - |
| Rat $Kp_{u,u}$ bolus, 5 minutes p.i. | >0.3 | 0.5 | 0.7 | - |

**Table S1.** ADME data for BIO-462, BIO-837, and BIO-687. Abbreviations: R (rat), H (Human), Cy (cynomolgus monkey), ER (efflux ratio), LM (liver microsomes).

#### Figure S1

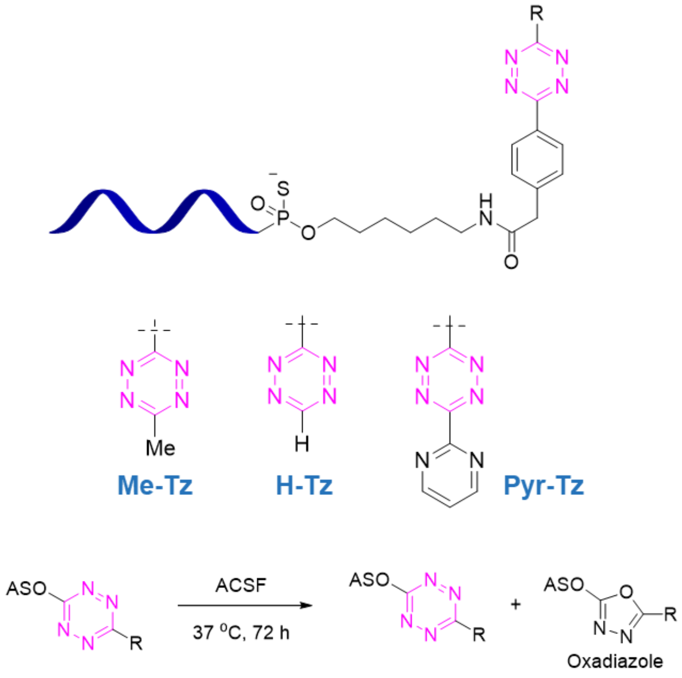

**Fig. S1.** Design of Malat1 ASO-tetrazine. (Top) C6-amine linker on the 5' end of the ASO; (bottom) three tested tetrazine substituents, left-to-right methyl-, proteo- , and pyridyl.

#### Figure S2

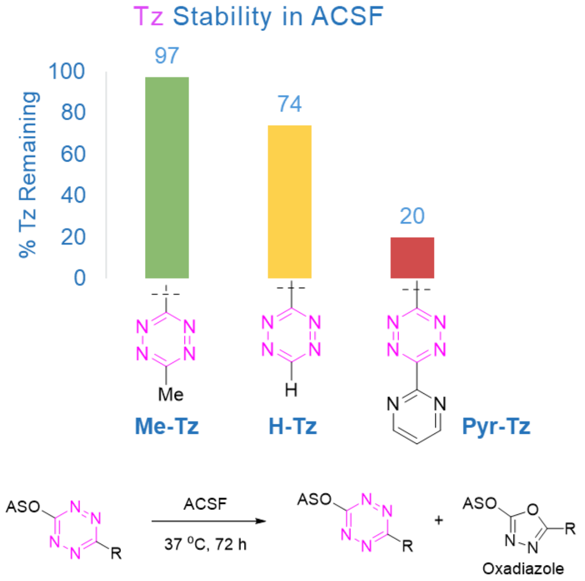

**Fig. S2.** Stability of three tested ASO-tetrazines in aCSF at 37 °C for 72 hours. (Bottom) initial stability conditions tested with observed oxidation product.

#### Figure S3

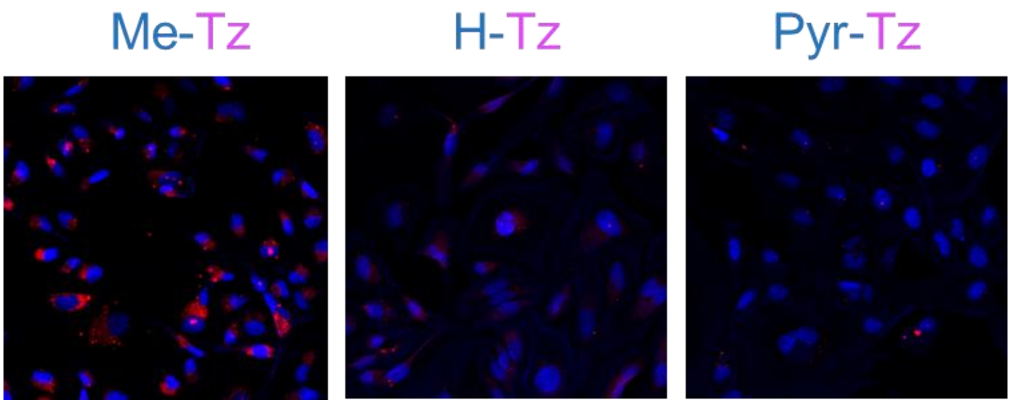

**Fig. S3.** In vitro uptake of three tested ASO-tetrazines in HeLa cells.

#### Figure S4

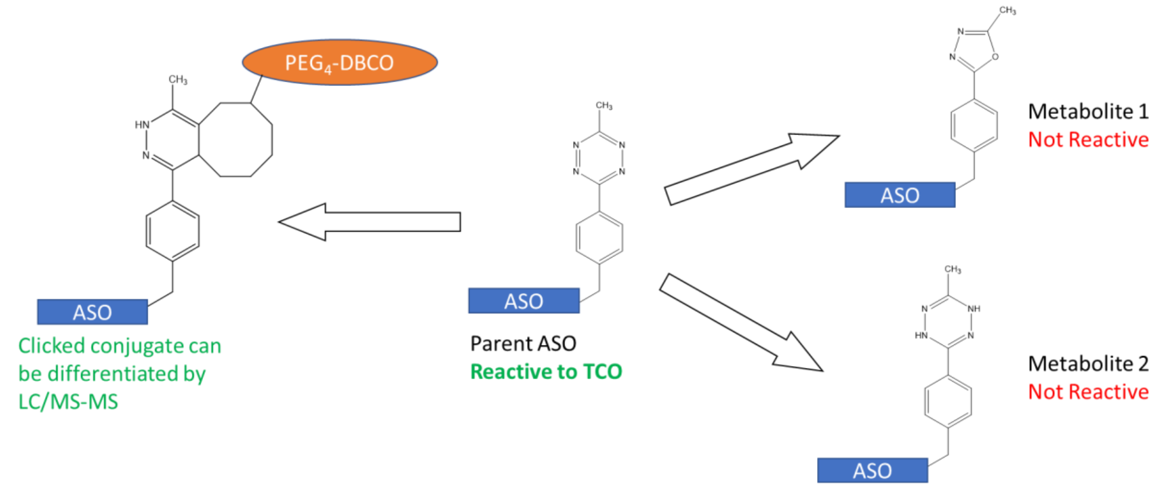

**Fig. S4.** Diagram illustrating how a DBCO-PEG$_4$-TCO probe is used to differentiate click-reactive ASO-MeTz from non-reactive metabolites using LC-MS.

### Figure S5

|  | Rat | Monkey | Human |
|---|---|---|---|
| Parent | 0 | 0 | 6.2 |
| M01 | 8.2 | 10.9 | <2 |
| M02 | 30.2 | 14.2 | <2 |
| M03 | 7.2 | 5.6 | 7.9 |
| M04 | 20.1 | 2.7 | 2.9 |
| M05 | 3.7 | 9.3 | 6.5 |
| M06 | 6.9 | <2 | <2 |
| M07 | 3.2 | 2.3 | <2 |
| M08 | 3 | 2.6 | <2 |
| M09 | 11.7 | 20.4 | 16.6 |
| M10 | <2 | 16.2 | <2 |
| M11 | <2 | 10.4 | <2 |
| M12 | <2 | <2 | 3.5 |
| M13 | <2 | <2 | 41.6 |
| M14 | ND | ND | 5.6 |

ND: not detected

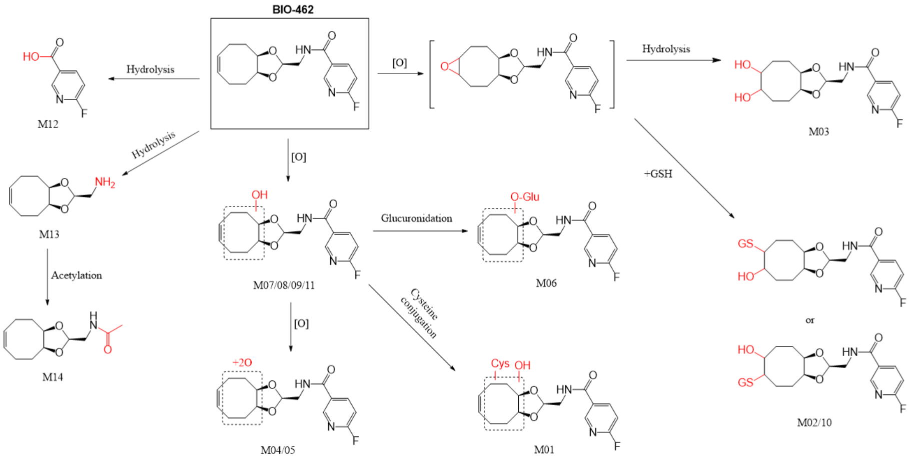

**Fig. S5.** Summary of results from metabolite identification study for BIO-462. Table at left shows percentage of each isolated metabolite from the different species of hepatocytes tested.

### Figure S6

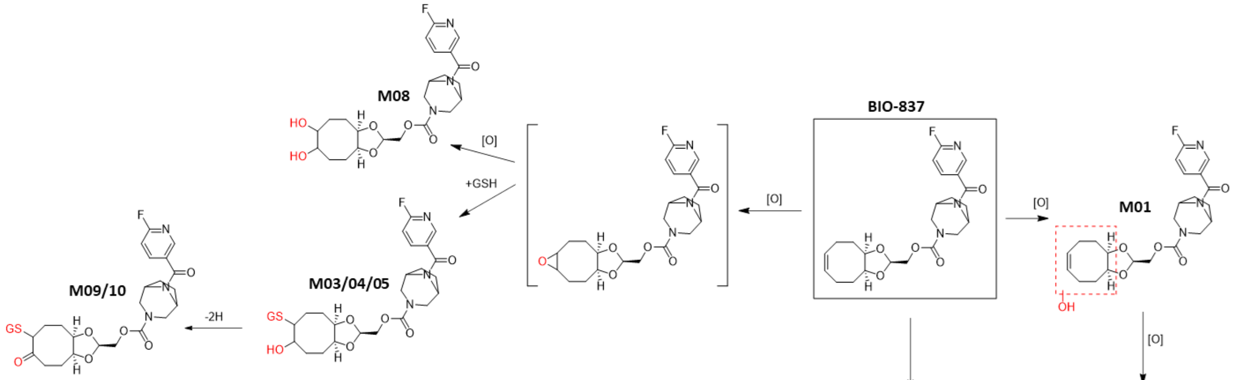

|  | Human | Rat | Monkey |
|---|---|---|---|
| Parent | 33 |  | 14 |
| M01 |  |  | 8 |
| M02 |  |  | 5 |
| M03 | 16 | 9 | 27 |
| M04 | 8 |  | 5 |
| M05 |  |  | 7 |
| M06 | 7 | 5 | 8 |
| M07 | 28 | 61 | 21 |
| M08 | 9 |  | 5 |
| M09 |  | 14 |  |
| M10 |  | 12 |  |

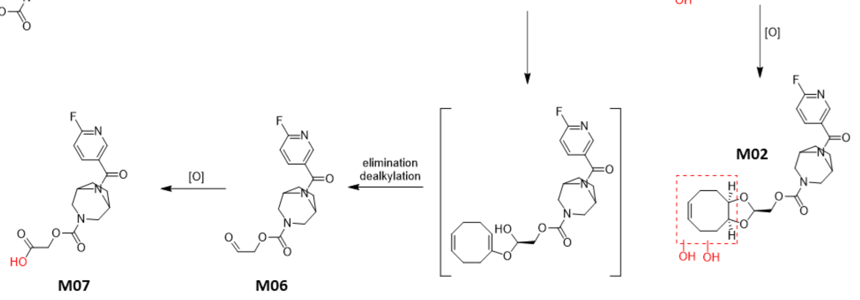

**Fig. S6.** Summary of results from metabolite identification study for BIO-837. Table at left shows percentage of each isolated metabolite from the different species of hepatocytes tested.

#### Figure S7

Semi-Prep HPLC chromatogram

Analytical HPLC chromatogram

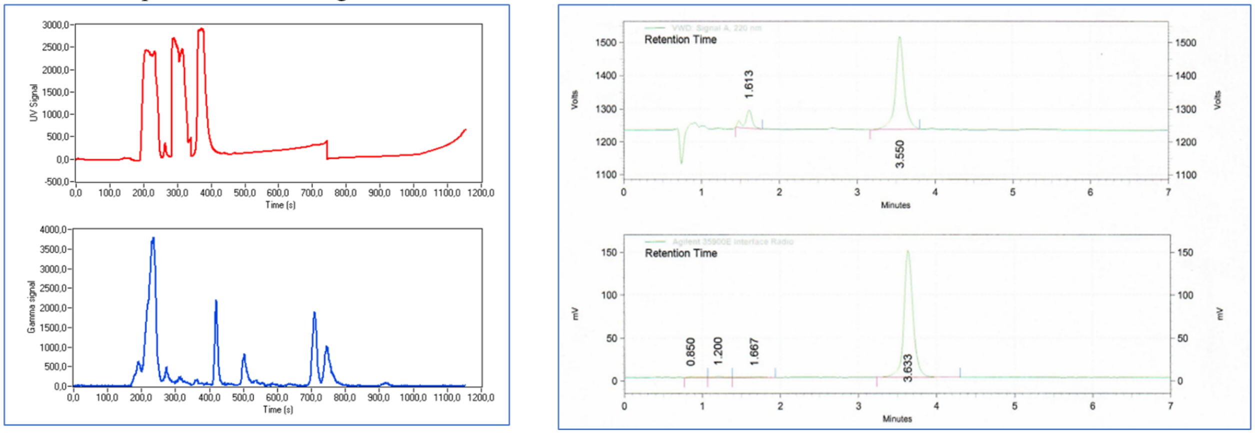

**Fig. S7.** Representative HPLC chromatograms from semi-preparative and analytical runs of the final product [$^{18}$F]BIO-687 acquired during radiosynthesis.

#### Figure S8

| Study Species | Synthesis # | RCY (n.d.c) | RCP at EOS | Molar activity (GBq/µmol) |
|---|---|---|---|---|
| Rat | 1 | 3% | 92% | 8.33 |
| Rat | 2 | 3.3% | 77% | 29.2 |
| Rat | 3 | 2% | 88% | 8.16 |
| Rat | 4 | 2.4% | 81% | 17.3 |
| Rat | 5 | 4.5% | 90% | 3.49 |
| Rat | 6 | 4.4% | 90% | 9.64 |
| NHP | 7 | 11% | >99% | 204 |
| NHP | 8 | 10% | >98% | 181 |
| NHP | 9 | 7% | >99% | 142 |

| | | | | |
|---|---|---|---|---|
| NHP | 10 | 12% | >99% | 203 |
| NHP | 11 | 2.5% | >98% | 257 |
| NHP | 12 | 2% | >96% | 75 |

**Fig. S8.** Summary of all radiolabeling syntheses of [$^{18}$F]BIO-687 (n.d.c. = non-decay corrected).

#### Figure S9

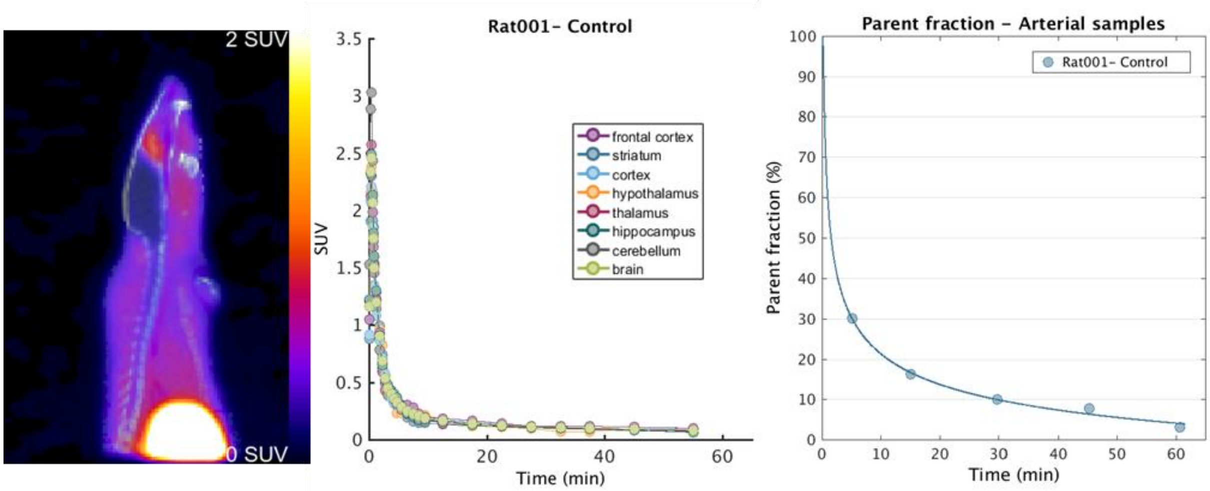

**Fig. S9.** Representative baseline PET/CT scan in naïve rat following i.v. dosing with [$^{18}$F]BIO-687; (left) maximum intensity projection (MIP) summing images from 0-60 minutes post-injection, (center) time-activity curves for select subregions and whole brain, (right) parent fraction of [$^{18}$F]BIO-687 from arterial blood samples taken at intervals throughout scan duration.

#### Figure S10

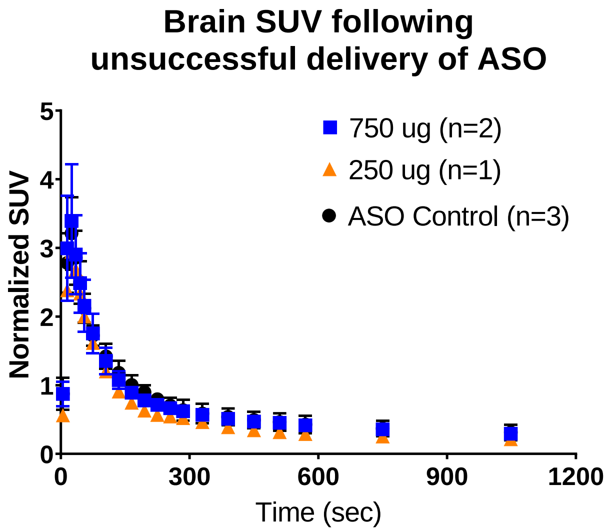

**Fig. S10.** Time-activity curves of whole brain in rats following i.t. dosing of ASO-MeTz and i.v. dosing with [$^{18}$F]BIO-687. It was determined during postmortem analysis that these rats in the 750 µg and 250 µg cohorts had levels of ASO BLQ in the brain and were excluded from analysis.

#### Figure S11

| | | Sample ID. | Frontal cortex | Striatum | Hippocampus | Thalamus+ Hypothalamus | Cerebellum | Cervical spinal cord | Thoracic spinal cord | Lumbar spinal cord | Liver | Kidney |
|---|---|---|---|---|---|---|---|---|---|---|---|---|
| High Dose (750 µg) | Rat 4 | MALAT ASO | 1820 | 591 | 2010 | 1200 | 1070 | 2170 | 2480 | 2180 | 508 | 3090 |
| | | MALAT-MeTz | 11500 | 2230 | 13900 | 7980 | 8700 | 31400 | 37900 | 26900 | 9060 | 32400 |
| | Rat 5 | MALAT ASO | 612 | 194 | 1560 | 1030 | 1270 | 1980 | 2590 | 2830 | 353 | 3030 |
| | | MALAT-MeTz | 2640 | 503 | 9490 | 6280 | 16800 | 31600 | 40800 | 42400 | 5660 | 27800 |
| | Rat 7 | MALAT ASO | BQL | BQL | BQL | BQL | BQL | BQL | 37.6 | 280 | 338 | 1770 |
| | | MALAT-MeTz | BQL | BQL | BQL | BQL | 44.9 | 92.9 | 308 | 6970 | 6440 | 19600 |
| | Rat 12 | MALAT ASO | BQL | BQL | BQL | BQL | BQL | BQL | 29.2 | 52.4 | 458 | 2430 |
| | | MALAT-MeTz | BQL | BQL | BQL | BQL | 52.3 | 112 | 236 | 514 | 8840 | 25100 |
| | Rat 19 | MALAT ASO | 1130 | 238 | 1230 | 767 | 1120 | 1710 | 2420 | 2430 | 392 | 4140 |
| | | MALAT-MeTz | 8650 | 904 | 9310 | 5270 | 12400 | 28300 | 49000 | 45000 | 10300 | 51400 |

| | | | | | | | | | | | | |
|---|---|---|---|---|---|---|---|---|---|---|---|---|
| Medium Dose (500 µg) | Rat 13 | MALAT ASO | 985 | 447 | 1010 | 863 | 851 | 1030 | 1720 | 820 | 311 | 1850 |
| | | MALAT-MeTz | 5680 | 2020 | 8140 | 6340 | 10700 | 11900 | 27900 | 10000 | 7420 | 20600 |
| | Rat 15 | MALAT ASO | 309 | 123 | 334 | 357 | 189 | 556 | 423 | 180 | 225 | 953 |
| | | MALAT-MeTz | 1450 | 374 | 1270 | 1670 | 1460 | 4270 | 3940 | 2030 | 4250 | 11600 |
| | Rat 18 | MALAT ASO | 612 | 157 | 609 | 801 | 712 | 1080 | 1510 | 1530 | 231 | 1660 |
| | | MALAT-MeTz | 3530 | 574 | 5310 | 7190 | 9370 | 13200 | 25000 | 24100 | 5000 | 20300 |
| | Rat 20 | MALAT ASO | 698 | 186 | 752 | 848 | 995 | 1290 | 1590 | 2020 | 260 | 1900 |
| | | MALAT-MeTz | 3330 | 624 | 4480 | 5550 | 13000 | 14400 | 22900 | 34500 | 6160 | 21000 |
| Low Dose (250 µg) | Rat 14 | MALAT ASO | BQL | BQL | BQL | BQL | BQL | BQL | BQL | BQL | 93.1 | 381 |
| | | MALAT-MeTz | BQL | BQL | BQL | 10.1 | BQL | 21.1 | 20 | 75.1 | 1570 | 3080 |
| | Rat 16 | MALAT ASO | 40.8 | 13.4 | 131 | 98.7 | 132 | 209 | 466 | 286 | 243 | 1140 |
| | | MALAT-MeTz | 104 | 47.4 | 360 | 371 | 763 | 2110 | 6280 | 3140 | 4770 | 11800 |
| | Rat 17 | MALAT ASO | 997 | 133 | 1040 | 624 | 560 | 737 | 566 | 341 | 151 | 1120 |
| | | MALAT-MeTz | 5170 | 404 | 6120 | 3380 | 6850 | 9470 | 6220 | 3770 | 3060 | 13400 |

**Fig. S11.** Results of LC/MS-MS analysis of Malat1 ASO-MeTz concentration in rat brain subregion homogenate. MALAT ASO denotes ASO in tissue that is not reactive to TCO-DBCO (presumed cleavage of MeTz linker from the ASO), while MALAT-MeTz denotes ASO that is reactive to TCO-DBCO probe.

#### Figure S12

| **Tissue Sample** | **Average** | **Std Dev** | **A04** | **A05** | **A07** | **A12** | **A19** | **A13** | **A15** | **A18** | **A20** | **A14** | **A16** | **A17** |
|---|---|---|---|---|---|---|---|---|---|---|---|---|---|---|
| Frontal cortex | 83.0% | 4.7% | 0.86 | 0.81 | - | - | 0.88 | 0.85 | 0.82 | 0.85 | 0.83 | - | 0.72 | 0.84 |
| Striatum | 77.4% | 2.8% | 0.79 | 0.72 | - | - | 0.79 | 0.82 | 0.75 | 0.79 | 0.77 | - | 0.78 | 0.75 |
| Hippocampus | 84.9% | 5.3% | 0.87 | 0.86 | - | - | 0.88 | 0.89 | 0.79 | 0.90 | 0.86 | - | 0.73 | 0.85 |
| Thalamus+ Hypothalamus | 85.6% | 3.3% | 0.87 | 0.86 | - | - | 0.87 | 0.88 | 0.82 | 0.90 | 0.87 | - | 0.79 | 0.84 |
| Cerebellum | 90.9% | 2.7% | 0.89 | 0.93 | - | - | 0.92 | 0.93 | 0.89 | 0.93 | 0.93 | - | 0.85 | 0.92 |
| Cervical spinal cord | 92.3% | 1.8% | 0.94 | 0.94 | - | - | 0.94 | 0.92 | 0.88 | 0.92 | 0.92 | - | 0.91 | 0.93 |
| Thoracic spinal cord | 92.6% | 2.2% | 0.94 | 0.94 | 0.89 | 0.89 | 0.95 | 0.94 | 0.90 | 0.94 | 0.94 | - | 0.93 | 0.92 |
| Lumbar spinal cord | 93.1% | 1.7% | 0.93 | 0.94 | 0.96 | 0.91 | 0.95 | 0.92 | 0.92 | 0.94 | 0.94 | - | 0.92 | 0.92 |
| Liver | 95.2% | 0.7% | 0.95 | 0.94 | 0.95 | 0.95 | 0.96 | 0.96 | 0.95 | 0.96 | 0.96 | 0.94 | 0.95 | 0.95 |

| | | | | | | | | | | | | | |
|---|---|---|---|---|---|---|---|---|---|---|---|---|---|
| Kidney | 91.5% | 1.0% | 0.91 | 0.90 | 0.92 | 0.91 | 0.93 | 0.92 | 0.92 | 0.92 | 0.92 | 0.89 | 0.91 | 0.92 |

**Fig. S12.** Ratio of reactive to non-reactive ASO-MeTz in postmortem rat tissues 24 hours after dosing i.t.

#### Figure S13

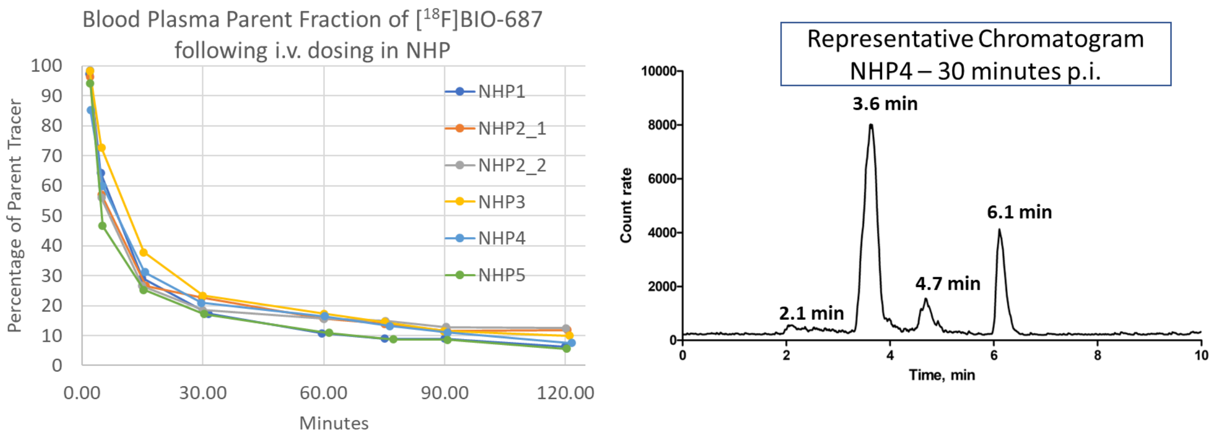

**Fig. S13.** Plasma parent fraction of [$^{18}$F]BIO-687 for each PET scan in NHP. Representative radioHPLC chromatogram from NHP4 30-minutes post-injection.

#### Figure S14

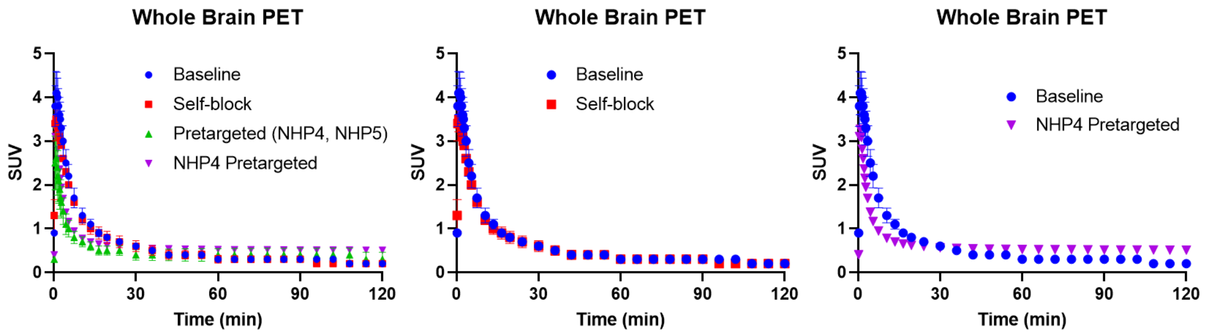

**Supplemental Fig. S14.** Time-activity curves for whole-brain PET ROIs in NHP.

### Figure S16

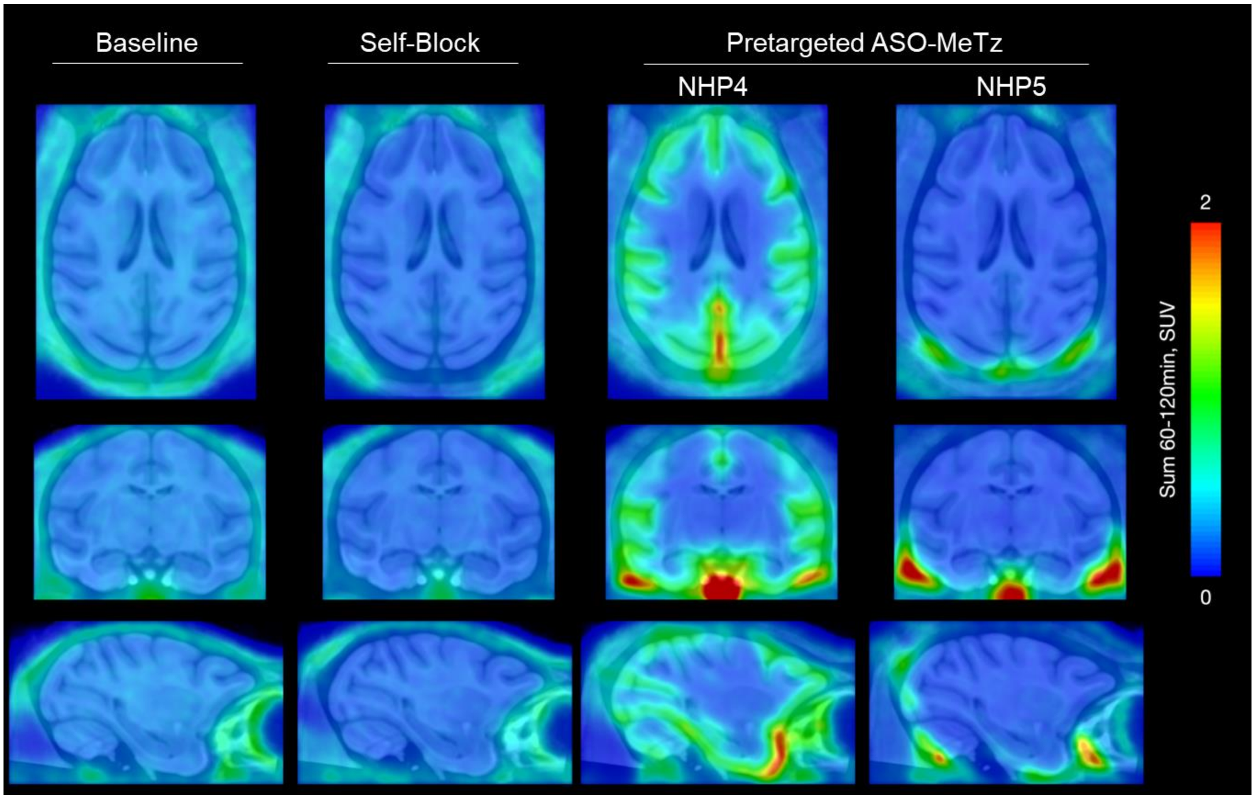

**Fig. S16.** Pretargeted PET imaging in cynomolgus monkeys. Left to right, baseline scan with [$^{18}$F]BIO-687, non-radioactive self-block with 1.0 mg/kg BIO-687, pretargeted with 20 mg Malat1 ASO-MeTz dosed IT followed 24 hours later with tracer (NHP4 and NHP5). Images are summed from 60-120 minutes post-injection. The static SUV images, in a range of 0-2, are overlaid on a standard template T1-weighted MRI of the cynomolgus monkey. The SUV images are smoothed at 2 mm FWHM.

### Chemistry

#### General procedure for the synthesis of Malat1-tetrazine analogues

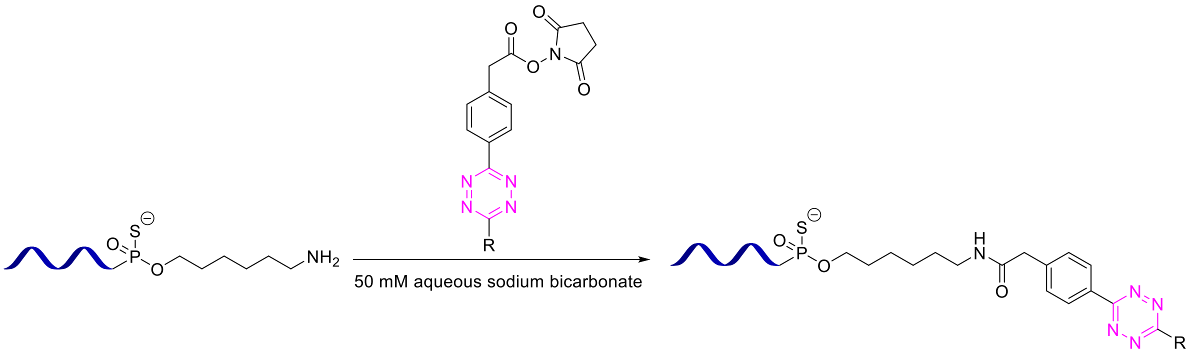

To an 8 mL vial containing Malat1 C6 amine (300 mL, 6.4 mM) in 50 mM aqueous sodium bicarbonate, a tetrazine-NHS ester (24 equiv.) was added as a solution in DMSO in three 330 uL portions over the course of 1 hour. The reaction was diluted with 8 mL of ethanol, and the mixture was cooled to 0 °C. The mixture was passed through a filter funnel, and the filter cake was washed with ethanol. The filter cake was taken up in water and loaded into an Amicon centrifugal filter (3 kDa cutoff), diluted to 5 mL, then spun down at 4000 RPM for 30 minutes. The mixture was concentrated to approximately 500 uL after spin filtering and was then diluted to 5 mL and spun down a second time. This process was performed a total of three times. The resulting solution was removed from the centrifugal filter, the filter was rinsed, and the combined material was lyophilized to afford the desired Malat1-Tz conjugate. Identity was confirmed by ICP-MS.

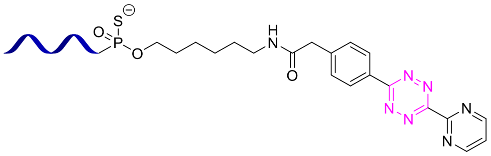

IPC-MS: m/z = 1904.3 (M-4H$^{+}$); z=-4,

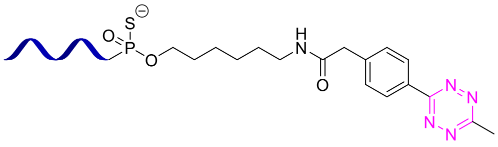

IPC-MS: m/z = 1888.3 (M-4H$^{+}$); z=-4

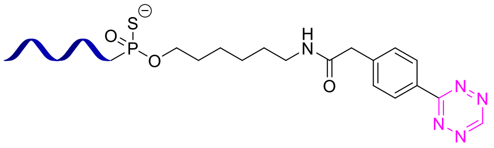

IPC-MS: m/z = 1884.8 (M-4H$^{+}$); z=-4

#### Synthesis of BIO-837 [Cis-cyclooctene Reference Standard]

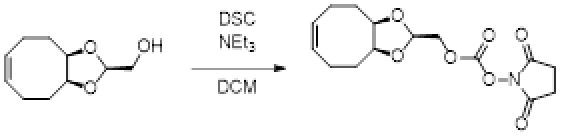

To a vial containing [(3aR,6Z,9aS)-3a,4,5,8,9,9a-hexahydrocycloocta[d][1,3]dioxol-2-yl]methanol (200 mg, 1.09 mmol), bis(2,5-dioxopyrrolidin-1-yl) carbonate (370.79 mg, 1.30 mmol, 90% purity), and DCM (2 mL) triethylamine (131.82 mg, 1.30 mmol, 181.57 µL) was added and the mixture was stirred at rt overnight. The following morning, the mixture was filtered over a pad of celite and concentrated. The crude residue was subjected to flash chromatography on silica (0-50% EtOAc in heptane), and after pooling and concentrating the appropriate fractions, [(3aR,6Z,9aS)-3a,4,5,8,9,9a-hexahydrocycloocta[d][1,3]dioxol-2-yl]methyl (2,5-dioxopyrrolidin-1-yl) carbonate (217.5 mg, 668.59 µmol, 61.59% yield) was obtained as a colorless oil.

$^{1}$H NMR (400 MHz, $CDCl_3$) δ ppm 5.57 - 5.66 (m, 2 H), 5.10 (t, J=3.3 Hz, 1 H), 4.37 (d, J=3.3 Hz, 2 H), 4.18 - 4.29 (m, 2 H), 2.80 - 2.88 (m, 4 H), 2.45 - 2.62 (m, 2 H), 1.95 - 2.22 (m, 6 H).

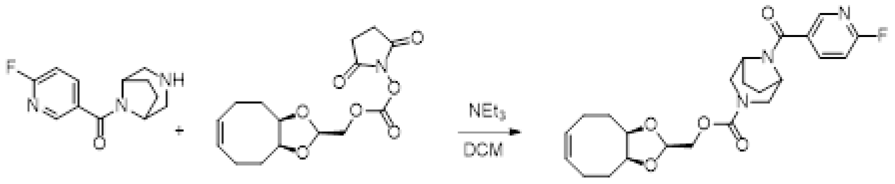

To a vial containing 3,8-diazabicyclo[3.2.1]octan-8-yl-(6-fluoro-3-pyridyl)methanone (128.47 mg, 368.88 µmol, TFA) and TEA (311.05 mg, 3.07 mmol, 428.45 uL) in DCM (2 mL), [(3aR,6Z,9aS)-3a,4,5,8,9,9a-hexahydrocycloocta[d][1,3]dioxol-2-yl]methyl (2,5-dioxopyrrolidin-1-yl) carbonate (100 mg, 307.40 µmol) was added and the mixture was stirred at rt for 3 hours. The volatiles were removed under reduced pressure and the crude residue was subjected to flash chromatography on silica (0-80% EtOAc in heptane). After pooling and concentrating the appropriate fractions, [(3aR,6Z,9aS)-3a,4,5,8,9,9a-hexahydrocycloocta[d][1,3]dioxol-2-yl]methyl 8-(6-fluoropyridine-3-carbonyl)-3,8-diazabicyclo[3.2.1]octane-3-carboxylate (119.7 mg, 268.70 µmol, 87.41% yield) was obtained as a colorless oil.

LCMS: m/z = 446 $(M+H)^+$

#### Synthesis of BIO-687 Trans-cyclooctene Reference Standard

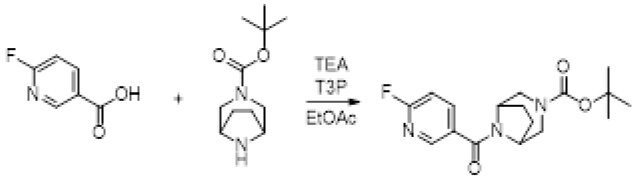

To a vial containing 6-fluoropyridine-3-carboxylic acid (200 mg, 1.42 mmol), tert-butyl 3,8-diazabicyclo[3.2.1]octane-3-carboxylate (361.09 mg, 1.70 mmol), TEA (717.15 mg, 7.09 mmol, 987.81 µL), and EtOAc (2 mL), T3P (2.71 g, 4.25 mmol, 2.53 mL, 50% purity) was added and the mixture was stirred at rt for 3 hours. The mixture was concentrated under reduced pressure and the resultant crude oil was subjected to flash chromatography on silica gel (0-50% EtOAc in heptane). After pooling and concentrating the appropriate fractions, tert-butyl 8-(6-fluoropyridine-3-carbonyl)-3,8-diazabicyclo[3.2.1]octane-3-carboxylate (465 mg, 1.39 mmol, 97.82% yield) was obtained as a colorless oil.

LCMS: m/z = 280 $(M+H\text{-isobutylene})^+$

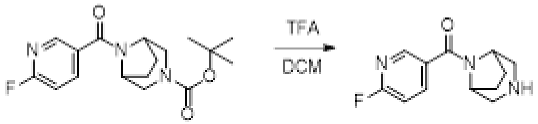

A vial containing tert-butyl 8-(6-fluoropyridine-3-carbonyl)-3,8-diazabicyclo[3.2.1]octane-3-carboxylate (460.00 mg, 1.37 mmol), DCM (5 mL), and TFA (2.98 g, 26.12 mmol, 2 mL) was stirred at rt overnight. In the morning, the mixture was concentrated under reduced pressure to afford a pale-yellow oil that was used without further purification.

LCMS: m/z = 236 $(M+H)^+$

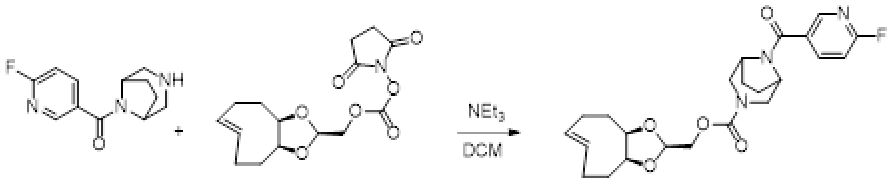

To a vial containing 3,8-diazabicyclo[3.2.1]octan-8-yl-(6-fluoro-3-pyridyl)methanone (43.39 mg, 184.44 µmol) and TEA (155.53 mg, 1.54 mmol, 214.22 uL) in DCM (1 mL), [(3aR,6E,9aS)-3a,4,5,8,9,9a-hexahydrocycloocta[d][1,3]dioxol-2-yl]methyl (2,5-dioxopyrrolidin-1-yl) carbonate[1] (50.00 mg, 153.70 µmol) was added and the mixture was stirred at rt for 1 hour. The mixture was concentrated under reduced pressure and the crude residue was subjected to flash chromatography on silica gel (0-80% EtOAc in heptane). After pooling and concentrating the appropriate fractions, [(3aR,6E,9aS)-3a,4,5,8,9,9a-hexahydrocycloocta[d][1,3]dioxol-2-yl]methyl 8-(6-fluoropyridine-3-carbonyl)-3,8-diazabicyclo[3.2.1]octane-3-carboxylate (36.5 mg, 81.93 µmol, 53.31% yield) was obtained as a colorless oil.

LCMS: m/z = 446 $(M+H)^+$

$^{1}H$ NMR (500 MHz, DMSO-d6) δ ppm 8.42 (d, J=2.4 Hz, 1 H) 8.14 (td, J=8.1, 2.1 Hz, 1 H) 7.29 (dd, J=8.5, 2.4 Hz, 1 H) 5.52 - 5.61 (m, 2 H) 4.84 (br s, 1 H) 4.68 (br s, 1 H) 3.60 - 4.16 (m, 8 H) 3.19 (br d, J=11.6 Hz, 1 H) 3.09 (br d, J=12.2 Hz, 1 H) 2.29 (dt, J=13.6, 6.9 Hz, 1 H) 2.03 - 2.21 (m, 3 H) 1.79 - 1.95 (m, 4 H) 1.54 - 1.69 (m, 4 H) 1.46 (br s, 1 H)

#### Synthesis of Compound 1 [TCO-precursor]

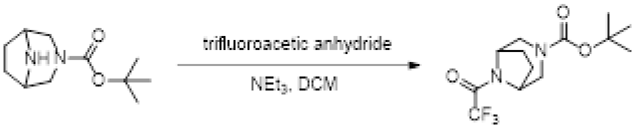

To a solution of tert-butyl 3,8-diazabicyclo[3.2.1]octane-3-carboxylate (2.9 g, 13.66 mmol) in DCM (25 mL) was added TEA (2.07 g, 20.49 mmol, 2.86 mL) and (2,2,2-trifluoroacetyl) 2,2,2-trifluoroacetate (3.16 g, 15.03 mmol, 2.09 mL) at 0 °C. The mixture was stirred at 0 °C to 25 °C for 1 hour.  .  Water (50 mL) was added and the mixture was extracted with DCM (20 mL x 3).  The organic layer was washed with brine and dried over Na2SO4.  The insolubles were filtered, the filtrate was concentrated under reduced pressure, and the thus obtained residue was purified by Combi Flash (EA in PE from 0 % to 9 %) to give tert-butyl 8-(2,2,2-trifluoroacetyl)-3,8-diazabicyclo[3.2.1]octane-3-carboxylate (3.66 g, 10.87 mmol, 79.59% yield, 91.58% purity) as yellow oil.

LCMS: m/z = 253 $(M+H\text{-isobutylene})^{+}$

$^{1}H$ NMR (500MHz, CHLOROFORM-d) δ = 4.73-4.69 (m, 1H), 4.42-4.37 (m, 1H), 4.02 (s, 1H), 3.86 (t, J = 13.0 Hz, 1H), 3.19-2.94 (m, 2H), 2.04-1.76 (m, 4H), 1.53-1.43 (m, 9H).

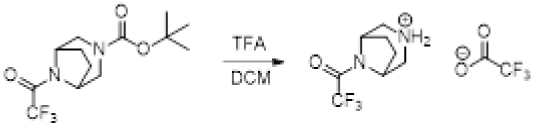

To a solution of tert-butyl 8-(2,2,2-trifluoroacetyl)-3,8-diazabicyclo[3.2.1]octane-3-carboxylate (3.66 g, 11.87 mmol) in DCM (20 mL) was added TFA (14.89 g, 130.59 mmol, 10 mL).  The mixture was stirred at 25 °C for 16 hours.  LCMS (BIIB-100615-329-P1A) showed the starting material was consumed completely and the desired MS was found.  The mixture was concentrated to give the trifluoroacetate salt of 1-(3,8-diazabicyclo[3.2.1]octan-8-yl)-2,2,2-trifluoro-ethanone (4.5 g, crude) as light brown oil that was used without further purification.

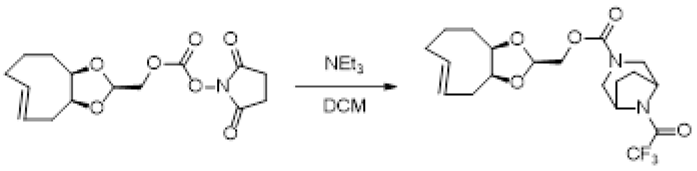

To a vial containing 1-(3,8-diazabicyclo[3.2.1]octan-8-yl)-2,2,2-trifluoro-ethanone (118.48 mg, 368.88 µmol, TFA) and NEt3 (311.05 mg, 3.07 mmol, 428.45 uL) in DCM (2 mL), [(3aR,6E,9aS)-3a,4,5,8,9,9a-hexahydrocycloocta[d][1,3]dioxol-2-yl]methyl (2,5-dioxopyrrolidin-1-yl) carbonate1 (100 mg, 307.40 µmol) was added and the mixture was stirred at rt for 1 hour. The volatiles were removed under reduced pressure and the mixture was subjected to flash chromatography on silica gel (0-50% EtOAc in heptane). After pooling and concentrating the appropriate fractions, [(3aR,6E,9aS)-3a,4,5,8,9,9a-hexahydrocycloocta[d][1,3]dioxol-2-yl]methyl 8-(2,2,2-trifluoroacetyl)-3,8-diazabicyclo[3.2.1]octane-3-carboxylate (55 mg, 131.45 µmol, 42.76% yield) was obtained as a pale yellow oil.

$^{1}$H NMR (500 MHz, methanol-d4) δ ppm 5.51 - 5.72 (m, 2 H) 4.88 - 4.94 (m, 1 H) 4.63 - 4.77 (m, 1 H) 4.40 - 4.57 (m, 1 H) 3.89 - 4.21 (m, 6 H) 3.05 - 3.23 (m, 2 H) 2.33 - 2.43 (m, 1 H) 2.21 - 2.30 (m, 1 H) 2.04 - 2.21 (m, 3 H) 1.46 - 2.00 (m, 7 H).

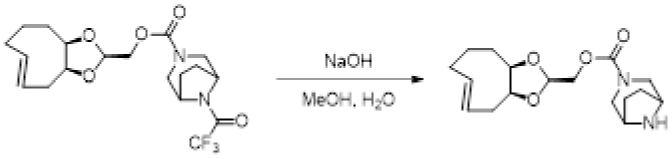

To a vial containing [(3aR,6E,9aS)-3a,4,5,8,9,9a-hexahydrocycloocta[d][1,3]dioxol-2-yl]methyl 8-(2,2,2-trifluoroacetyl)-3,8-diazabicyclo[3.2.1]octane-3-carboxylate (55 mg, 131.45 µmol) in MeOH (3 mL), NaOH (52.58 mg, 1.31 mmol) in water (1 mL) was added, and the mixture was heated to 60 °C for 1 hour. The mixture was cooled to rt and the volatile organics were removed under reduced pressure. The mixture was diluted with water and extracted with three portions of DCM. The combined organics were passed over a plug of magnesium sulfate and concentrated to afford a colorless film that was used without further purification.

$^{1}$H NMR (500 MHz, acetonitrile-d3) δ ppm 5.50 - 5.70 (m, 2 H), 5.14 (t, J=4.0 Hz, 1 H), 4.84 (t, J=3.7 Hz, 1 H), 3.86 - 4.11 (m, 4 H), 3.55 - 3.70 (m, 2 H), 3.36 (br d, J=13.4 Hz, 2 H), 2.92 (br dd, J=42.7, 12.2 Hz, 2 H), 2.29 - 2.39 (m, 1 H), 2.00 - 2.26 (m, 4 H), 1.81 - 1.92 (m, 1 H), 1.45 - 1.74 (m, 7 H).

#### Radiochemistry

The fully optimized methods used at Karolinska for the NHP study are reported here, with similar methods used at Invicro for the rodent study. Fluorine-18 fluoride ($[^{18}F]F^-$) was produced from a GEMS PET trace Cyclotron using 16.4 MeV protons via the $^{18}O(p,n)^{18}F$ reaction on $^{18}O$ enriched water ($[^{18}O]H_2O$) and $[^{18}F]F$- was isolated from $[^{18}O]H_2O$ on a preconditioned Sep-Pak® Light Accell™ Plus QMA Carbonate anion exchange cartridge and subsequently eluted from the cartridge with a solution of Tetrabutylammonium hydrogen carbonate solution (0.075 M, 0.5 mL) and acetonitrile (0.5 mL) to a reaction vessel (reactor 1, 4.5 mL). The solvents were evaporated at 150 °C for 10 minutes under continuous nitrogen flow (70 mL/min) to form a dry complex of $[^{18}F]F^-/K_2CO_3/K_{2.2.2}$ and the residue was cooled to room temperature (RT).

The PyTFP precursor (10 mg in 1.0 mL of t-Butanol: anhydrous Acetonitrile (8:2)) was added to the dry $[^{18}F]$fluoride. The closed reaction vessel was heated at 80 °C for 10 minutes. The reaction mixture was cooled to RT and was diluted with water (10 mL) before loading on to a pre-conditioned (10 mL acetonitrile followed by 10 mL water) Oasis® MCX cartridge. The cartridge was washed with Acetonitrile: water (1:9, 5 mL) and dried under $N_2$-flow for 5 minutes. Desired product $[^{18}F]$FPyTFP was eluted off the cartridge using acetonitrile (1.5 mL) to a second reaction vessel (reactor 2, 4.5 mL) containing Compound 1 (10 mg in 0.5 mL acetonitrile). The reaction mixture was left at 37 °C for 10 minutes. After the reaction, the mixture was diluted with water (2 mL) before injection onto the high-performance liquid chromatography (HPLC) column for purification. A semi-preparative reverse ACE 5 C18-HL column (C18, 7.8 Ø × 250 mm, 10 µm) was used for the purification. The column outlet was connected to an UV absorbance detector (λ=254 nm) in series with a GM-tube for radioactivity detection. Elution was performed with a mobile phase acetonitrile/water (35/65) at a flow rate of 6 mL/min. A radioactive fraction corresponding the pure $[^{18}F]$BIO-687 ($t_R$=28-30 min) was collected and diluted with sterile water (40 mL). The resulting mixture was loaded on to a pre-conditioned SepPak tC18 plus cartridge. The cartridge was washed with water (10 mL) and the isolated product, $[^{18}F]$BIO-687, was eluted with 1 mL of ethanol in to a sterile vial containing a mixture of tocoferol (50 mg), PEG (3 mL), and PBS (7 mL). The final product was filtered through a sterile filter (0.22 µm; Millipore, Bedford, MA), yielding a sterile and pyrogen free solution of [$^{18}$F]BIO-687.

The radiochemical purity, identity and stability of [$^{18}$F]BIO-687 was determined by analytical HPLC system which included a XBridge RP column (C18, 5 µm, 4.6 x 150 mm particle size), Merck-Hitatchi L-7100 Pump, L-7400 UV detector and GM-tube for radioactivity detection (VWR International) (Figure SU). The mobile phase $CH_3CN$/TFA (0.1%) with a gradient HPLC method (15-90% in 10 min) and flow rate of 1.5 mL/min was used to elute the product. The effluent was monitored with an UV absorbance detector (ƛ = 254 nm) coupled to a radioactive detector (b-flow, Beckman, Fullerton, CA). The identity of fluorine-18 labelled compounds was confirmed by using HPLC with the co-injection of the corresponding authentic non-radioactive reference standard.

The molar activity of the final product was measured by analytical HPLC which included a XBridge RP column (C18, 5 µm, 4.6 x 150 mm particle size) using mobile phase $CH_3CN$/TFA (0.1%) (0.05 M) with a gradient HPLC method (10-90% in 10 min) and flow rate of 2 mL/min. MA was calibrated for UV absorbance (ƛ = 254 nm) response per mass of ligand and calculated as the radioactivity of the radioligand (GBq) divided by the amount of the associated carrier substance (µmol).

### In Vitro Characterization of BIO-687

#### Brain and Plasma Protein Binding Assay

Compounds are prepared at 1 µM in either plasma or brain protein (diluted 1 part brain homogenate to 7 parts 1x PBS). Compounds are then incubated at 37°C and 5% $CO_2$ for 4 h. A 50 µL aliquot is then taken out and precipitated in 250 µL 1:1 ACN/methanol. Samples are centrifuged for 10 min at 3220 x g. an aliquot is then removed and diluted to 90:10 H2O/ACN + 0.1% formic acid to be run on HPLC.

#### Efflux Transporter (MDCK-MDR1/BCRP) Assay

Madin-Darby Canine Kidney cells (MDCK) that were singly transfected to express the mock empty vector (wild-type, WT), or human P-glycorptein (P-gp, MDR1; NIH cells licensed from Absorption Systems), or human breast cancer resistance protein (BCRP) efflux transporter were (acquired from Solvo Biotechnology, Szeged, Hungary). Cells were maintained in Dulbecco's Modified Eagle Medium (DMEM) with 10% fetal bovine serum (FBS), 1% Penicillin/Streptomycin (PEST) in a humidified incubator at 37°C with 5% $CO_2$. The culture medium was changed three times weekly, and cell growth was observed by light microscopy. The MDCK-MDR1, -BCRP or -WT cell lines were seeded into 96-well Corning transwell plates (cat #3392) for 4 or 5 d at 0.25 or 0.4 million cells/mL, respectively. Medium was changed once to twice per week prior to analysis.

Bidirectional flux was measured in triplicate wells in-house. Aliquots of test article DMSO stock were added to Hanks Balanced Salt Solution (HBSS) containing 4-(2-hydroxyethyl)-1-piperazineethanesulfonic acid (HEPES), at pH 7.4; the nominal concentration was 1.00 µM. The in-house protocol utilized 1% bovine serum albumin (BSA) in the receiver wells to negate non-specific binding. Monolayer confluence was monitored pre and post 2-hr incubation by trans epithelial electronic resistance (TEER) measurement. Monolayer confluence and active efflux were confirmed with commercially available compounds.

The samples were withdrawn at pre-selected time points (t=0 and 120 min) from the receiver and donor compartments and matrix-matched with either an equal volume of dH2O or 1% BSA in buffer and crashed with acetonitrile containing internal standard (IS). Samples were injected onto a high throughput liquid chromatography instrument (RapidFire, Agilent) and triple-quadrupole mass spectrometer (API5500, Sciex). The peak area ratio (PAR) of test article/IS were used for all calculations.

The impact of endogenous uptake and efflux transporters in the Solvo cell line was assessed by the parallel use of mock (WT) cells in each assay. The rationale for this inclusion is that the same scaffold of MDCK cells used in the transfection process would also display any similar uptake or efflux not directly attributed to the transfected protein, BCRP.

Monolayer confluence and active efflux were confirmed with commercially available compounds such as the low permeability compound, bestatin or atenolol; the Pgp substrate, loperamide; or the BCRP substrates, daidzein, cladribine, and PhIP.

Apparent permeability can be calculated using the equation: $Papp = (dQ / dt) / (C_0 \times A)$ where dQ/dt is the accumulation rate by volume (µmol/L.min), defined as the slope obtained from linear regression of test article transport amount at a fixed time. $C_0$ is the initial concentration donor side (µmol/L). A is the surface area of the filters or inserts (0.143 $cm^2$). The efflux ratio compares the permeabilities obtained in the basolateral-to-apical and opposite direct:

Efflux Ratio = Papp b-a/ Papp a-b

The efflux ratios require modification to take into consideration activity in the wild-type or control experiment: Corrected flux ratio = Efflux ratio BCRP / Efflux ratio WT.

##### Metabolite Identification for BIO-837 and BIO-462

*In vitro* metabolite profiling was performed in human, rat, and cynomolgus monkey hepatocytes (Lonza PN HUCS50P and RSCS01, Sekisui Xenotech PN PPCH2000). BIO-837 or BIO-462 (10 µM) were incubated in DMEM containing 1 million cells/mL for 1 hour (human) or 0.5 hours (rat and monkey) at 37°C under 5% $CO_2$. The reaction was stopped using a 1:1 addition of ice-cold acetonitrile. After centrifugation, the supernatant was diluted 1:3 with water for analysis. LC-MS was performed using a Waters UPLC and a Sciex 5600 Triple-TOF MS, with a Waters Acquity HSS T3 column (50 mm x 2.1 µm, 1.8 µm particle size) at 50˚C with a 20 µL injection volume. A linear gradient of 0.1% formic acid in water (A) and 0.1% formic acid in acetonitrile (B) was used from 5 - 55% B over 5 min, at 0.45 mL/min, followed by a wash of 90% B. HCD collision energy of 35 was used, and data was processed using Sciex PeakView and MetabolitePilot software.

#### In Vitro and Ex Vivo Characterization of Malat1 ASO-MeTz

##### Quantification of Malat1 ASO-MeTz in Tissue

Individual stock solution of MALAT1 ASO and MALAT1 ASO-MeTz were prepared at the concentration of 1 mg/mL. Standard and quality control (QC) working solutions were prepared by dilutions of stock solutions using ASO diluent (25 mM HFIP, 15 mM DMCHA, 100 µM EDTA, and 0.05 % rat plasma in water: ACN 90:10% v/v). Working solutions were spiked into blank matrix to yield calibration standards within the range of 1 – 1000 ng/mL.

Rat brain samples were weighed and homogenized in 9-fold (v/w) of lysis-loading buffer using a MP Biomedicals FastPrep-24 homogenizer, yielding 10-fold diluted brain homogenates. An aliquot of 200 µL calibration standards, QC samples and brain samples were added to the KingFisher 96 deep-well plate. 15 µL of probe-conjugated magnetic bead suspension was added to each sample to capture MALAT1 ASO and MALAT1 ASO-MeTz. The beads were transferred and sequentially washed three times. Finally, the beads were transferred to a pre-heated elution plate containing 200 µL ASO diluent with 25 ng/mL internal standard and rigorously agitated for 15 min at 90 °C to generate the hybridization extract for LC-MS/MS analysis. For the analysis of reactive MALAT1 ASO-MeTz, 10 µL of 500 µM TCO-PEG4-DBCO was added to each well to conjugate with MALAT1 ASO-MeTz. After incubation at room temperature for 120 hours, the samples were analyzed by LC-MS/MS.

An ExionLC AD UHPLC system with a Clarity 1.7 µm Oligo-XT 100 Å 50 × 2.1 mm column was used for the chromatographic separation of MALAT1 ASO, MALAT1 ASO-MeTz, MALAT1 ASO-MeTz-TCO-PEG4-DBCO and internal standard. The initial condition was 2% of mobile phase B, 98% of mobile phase A at 0.45 mL/min and 100% of mobile phase C at 0.05 mL/min. The initial condition was held for 0.2 min, the separation gradient ramped to 20% B within 5.5 min, while mobile phase C kept at 100% at 0.05 mL/min. After the gradient, the mobile phase was set to the initial condition and equilibrated for 1 min. Column temperature was set to 60 °C. The injection volume was 10 µL. MS/MS detection was conducted using a Sciex QTRAP 6500+ mass spectrometer equipped with an electrospray ionization (ESI) source at negative ion mode using multiple reaction monitoring (MRM). The optimized ion source parameters included curtain gas at 40, collision gas at high, ionspray voltage at -3500 V, temperature at 550 °C, ion source gas 1 at 60, ion source gas 2 at 60. MRM transitions were 793.1 to 95 for Malat1 ASO, 754.7 to 95 for Malat1 ASO-MeTz, 819.1 to 95 for Malat1 ASO-MeTz- TCO-PEG4-DBCO, and 879.5 to 95 for the internal standard.

#### Cell Uptake and Immunofluorescence Staining of Malat1 ASO-MeTz

The following protocol was adapted from Cook, et al. 2022 (4). Summarized in brief, HeLa cells were cultured in Eagle's Minimum Essential Medium (EMEM) with 10% fetal bovine serum, and incubated at 37 °C, 5% $CO_2$.

Cells were cultured for 24 hours in a 4-well cell culture slide (ThermoFisher) and stock solution of Malat1 ASO-MeTz in PBS was added to a final concentration of 5 µM in the cell media. The cells were incubated again for 24 h. Cells were then fixed with 4% PFA solution in PBS for 15 min at room temperature (RT) and permeabilized using a 0.1% Triton X-100 solution prior to staining. Primary antibodies were diluted in PBST (EEA1 at 1:100; LAMP1 at 1:200) and each on added to a single well (100 µL) and incubated for 45 min at 37 °C. The secondary antibody (Goat anti-rabbit IgG-FITC) was diluted 1:60 in PBST-BSA. Also to this solution was added TCO-Cy5 (Click Chemistry Tools #1089) to a final concentration of 1 µM. 100 µL of this solution was added to each well and incubated in the dark at RT for 1 h. 200 µL of 1x stock solution of Phalloidin-AF568 was added to each well and incubated for 20 min at RT. Wells were washed in PBS and then mounted using 30 µL of Prolong Gold + DAPI and allowed to cure overnight at RT in the dark.

Confocal imaging data was acquired using a fully motorized Zeiss Axio Observer Z1 (Carl Zeiss, Jena, Germany) inverted imaging system using a spinning disk confocal scanner unit CSU-W1 (Yokogawa), equipped with a 63x objective lens and two Hamamatsu ORCA-Flash4.0 v2 sCMOS cameras for 2 channel simultaneous acquisition. Solid-state lasers (405, 488, 561, 647 and 725 nm) were coupled to the spinning head through a fiber optic. A piezo PZ-2150 XYZ motorized stage was used to acquire 3-D stacks. Slidebook (Intelligent Imaging Innovations, Dencer, USA) was used to acquire, view, scale, process, and export images for publication and for image analysis.

#### Ex Vivo Autoradiography and Immunofluorescence of Malat1 ASO-MeTz

The Malat1 ASO-MeTz (500 µg in 30 µL aCSF, 40 µL flush) was dosed i.t. in naïve female Sprague-Dawley rats (n=4). Rats were allowed to recover for 24 hours, after which they were euthanized by $CO_2$ inhalation and brains removed and frozen on crushed dry ice. They were then sectioned to 10 micron slices and mounted on glass slides.

Slides were then air dried for 10 minutes before fixing in 4% paraformaldehyde (PFA) for 10 minutes. Samples were then blocked and permeabilized for 30 min in 1x PBS with 10% normal goat serum, 1% BSA, and 0.1% Triton X-100. Slides were incubated with anti-ASO primary antibody (generously provided by Ionis Pharmaceuticals) at a 1:3000 dilution in blocking buffer for 1 hour at room temperature. After washing with PBS, slides were incubated in a secondary cocktail of Alexa Fluor 488 Goat anti-Rabbit IgG H+L (Invitrogen Life Technologies A11008 2.0 mg/mL) at 3.0 µg/mL concentration and TCO-Cy5 1.0 µM concentration for 30 minutes at room temperature. Finally, slides were mounted with Prolong Diamond and coverslip. Confocal imaging was performed as above.

Sections near to those used for fluorescence staining were set aside for autoradiography and not fixed in PFA. These sections were then incubated with a solution of [$^{18}$F]BIO-687 (5 nM) for 15 min before being thoroughly washed with saline. Blocking solutions included 5 µM non-radioactive BIO-687 in addition to the radioactive tracer.

#### Longitudinal Distribution and Stability of Malat1 ASO-MeTz in Rat Brain

The Malat1 ASO-MeTz (500 µg in 30 µL aCSF, 40 µL flush) was dosed i.t. in naïve female Sprague-Dawley rats (n=10). Rats were allowed to recover for 24, 48, 96, 168, or 336 hours, after which they were euthanized by $CO_2$ inhalation and brains removed, bisected into left and right hemispheres, and frozen on crushed dry ice. Right hemispheres were then sectioned to 10 micron slices and mounted on glass slides, while left hemispheres were homogenized and analyzed by LC-MS/MS. Brain sections were stained with anti-ASO antibody and TCO-Cy5 as described above.

### In Vivo Studies in Rat and Non-Human Primates

Imaging and in vivo studies in rats and non-human primates were performed at Biogen, Invicro, and the Karolinska Institutet. Studies were carried out with oversite from institutional animal care and use committees in accordance with all local and regional laws.

#### Imaging tracer kinetics in naïve rat

Naïve male Sprague Dawley rats (N = 2, 273 ± 3 g) were placed under terminal anesthesia (isoflurane ca. 1.5 – 3%, 1 L oxygen·$min^{-1}$) and cannulae placed in an artery and a vein. Body temperature was monitored and maintained, and the respiration rate was monitored throughout the experiment. A dynamic PET scan of 60 minutes was carried out on each of the rats after IV injection of [$^{18}$F]BIO-687 (11.41 ± 1.92 MBq, 1.26 ± 0.78 µg), with a field of view focused on the brain and upper body. Arterial blood samples were collected continuously for the first 1 – 2 minutes, followed by discrete samples at 5, 15, 30, 45 and 60 minutes and metabolite analysis was carried out on the extracted plasma to generate the tracer input function for kinetics analysis. The rats were euthanized at the end of the scans.

Regions of interest were defined over 8 brain regions (frontal cortex, striatum, cortex, hypothalamus, thalamus, hippocampus, cerebellum, and whole brain) to generate the activity curves (TACs).

[1]H NMR

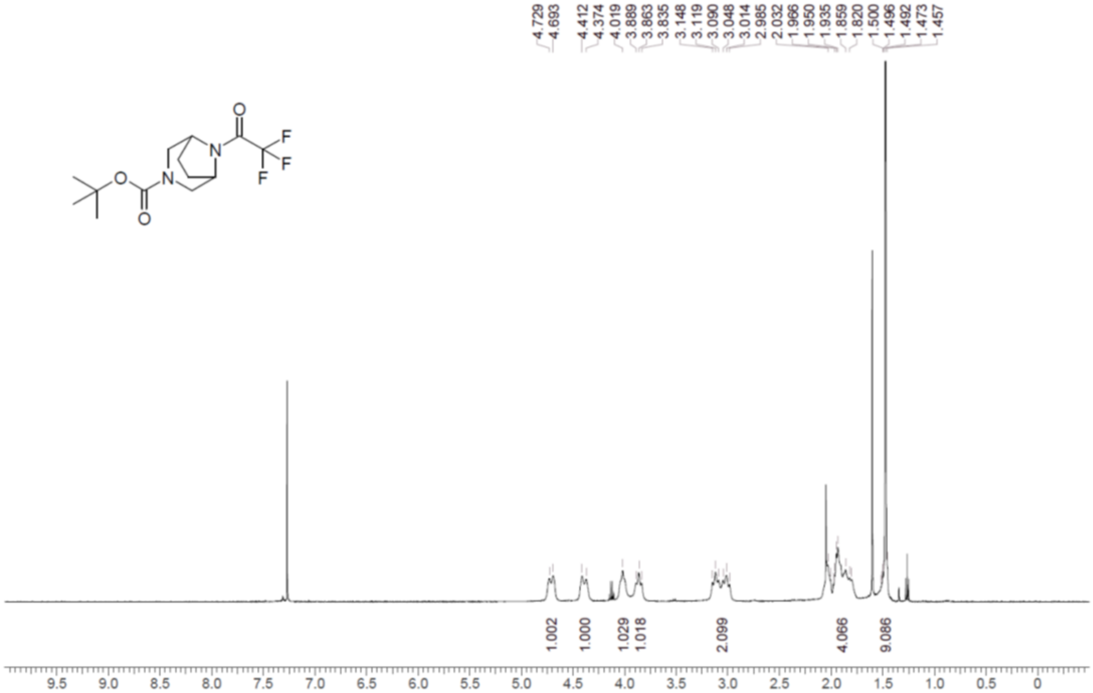
4.729
4.693
4.412
4.374
4.019
3.889
3.863
3.835
3.148
3.119
3.090
3.048
3.014
2.985
2.032
1.966
1.950
1.935
1.859
1.820
1.500
1.496
1.492
1.473
1.457
1.002
1.000
1.029
1.018
2.099
4.066
9.086
9.5
9.0
8.5
8.0
7.5
7.0
6.5
6.0
5.5
5.0
4.5
4.0
3.5
3.0
2.5
2.0
1.5
1.0
0.5
0
